# Lipid Metabolism and Contact Site Homologs Are Enriched at the Chloroplast Envelope-Thylakoid Interface

**DOI:** 10.1101/2025.06.20.660776

**Authors:** Evan W. LaBrant, Cailin N. Smith, Alondra D. Torres-Gerena, Joslin Ishimwe, Fan Huang, Allan Tullis, Lauren Litterer, Bhoomi Modi, Michael J. Naldrett, Bara Altartouri, Rebecca L. Roston

**Affiliations:** Department of Biochemistry, University of Nebraska-Lincoln, USA; Center for Plant Science Innovation, University of Nebraska-Lincoln, USA; Proteomics and Metabolomics Facility, Center for Biotechnology, University of Nebraska-Lincoln, USA; Morrison Microscopy Core Research Facility, Center for Biotechnology, University of Nebraska-Lincoln, USA; *Author for correspondence: (R.L.R.) N123 Beadle Center, 1901 Vine St., University of Nebraska-Lincoln Lincoln, NE 68516, USA

**Keywords:** Chloroplasts, Membrane Contact Sites, Thylakoid, Lipid Transport, Proteomics, Membrane Architecture

## Abstract

Biogenesis and maintenance of the photosynthetic thylakoid membrane requires transport of lipids from their site of synthesis in the chloroplast envelopes to their destination in the thylakoid. While vesicle trafficking is likely involved, we hypothesized a complementary mechanism involving direct membrane interactions. Using domain homology and proteomic profiling of chloroplast membrane fractions, we identified candidate lipid transport proteins enriched in a distinct, intermediate-density membrane population. This fraction was enriched in lipid metabolic enzymes and proteins homologous to known membrane organization factors. Several candidates, including TVP38 FAMILY PROTEIN (TVPFP), PLASMA MEMBRANE FUSION PROTEIN (PMFP), and LETM1-LIKE, localized to discrete subdomains within chloroplasts. Loss-of-function *tvpfp* or *pmfp* mutants exhibited altered chloroplast ultrastructure, including changes in thylakoid-envelope proximity, supporting their roles in maintaining membrane architecture. These findings identify a specialized chloroplast membrane region enriched in lipid-related functions and offer a foundation for elucidating the molecular architecture of these regions.

## INTRODUCTION

Chloroplasts are the sites of photosynthesis in plants and algae, containing a complex internal system of membranes called thylakoids. These membranes host the molecular machinery for light capture and energy conversion, embedded in a specialized lipid matrix that is conserved across all photosynthetic organisms (Petroutsos et al., 2014; Sato and Awai, 2016). The predominant lipids within thylakoids are sugar-lipids, in which monogalactosyldiacylglycerol and digalactosyldiacylglycerol are the most abundant, followed by sulfoquinovosyldiacylglycerol and a specialized phosphatidylglycerol (Boudière et al., 2014; Douce and Joyard, 1996). This specific lipid composition supports the assembly and maintenance of photosynthetic protein complexes (Gounaris et al., 1983; Mazur et al., 2020; Páli et al., 2003) and allows the tightly folded ultrastructure of mature thylakoid membranes (Fujii et al., 2019; Yoshihara et al., 2021).

Thylakoid maturation requires the coordinated integration of lipid, protein, and pigment synthesis to build functional photosynthetic membranes (Cackett et al., 2022; Jarvis and López-Juez, 2013). Many proteins necessary for thylakoid biogenesis have been identified, among them the multiple sets of protein import machinery (Zhu et al., 2022), retrograde signaling systems (Loudya et al., 2024), and glycerolipid biosynthetic enzymes (Cook et al., 2021). Because lipid synthesis occurs exclusively in the chloroplast envelopes, thylakoid membranes rely entirely on lipid transport from these sites (Hölzl and Dörmann, 2019). Proposed mechanisms for this transport include vesicle-mediated trafficking, membrane contact sites (MCSs), and inner envelope invaginations (LaBrant et al., 2018; Ostermeier et al., 2024).

During chloroplast development, inner envelope invaginations initiate early thylakoid membranes (Charuvi et al., 2012; Liang et al., 2018), but these structures are absent in mature plastids (Shimoni et al., 2005), pointing to additional mechanisms for thylakoid expansion. One such mechanism involves stromal vesicles, which have been observed by transmission electron microscopy (TEM; Lindquist et al., 2016) and are disrupted by loss of CHLOROPLAST-LOCALIZED SEC14-LIKE PROTEIN (CPSFL1) (García-Cerdán et al., 2020; Hertle et al., 2020; Kim et al., 2022). These vesicles are absent in *cpsfl1* even under conditions that normally enhance their production (Morré et al., 1991), and thylakoid formation is reduced but not eliminated (Hertle et al., 2020), implying the presence of another mechanism. A second vesicle-related protein, VESICLE-INDUCING PROTEIN IN PLASTIDS 1 (VIPP1/IM30) has also been implicated in the formation of stromal vesicles, although this is debated and it has additional roles in envelope membrane maintenance (Junglas et al., 2025; Zhang et al., 2012).

MCSs represent a non-vesicular alternative to lipid transfer. They are close membrane interfaces (∼20–50 nm) functionally specialized for lipid exchange and other shared chemistry functions (Lahiri et al., 2015; Quon and Beh, 2015). MCSs mediate lipid transfer between the ER and chloroplasts (Huercano et al., 2025) and from chloroplasts to mitochondria (Jouhet et al., 2024). Within the chloroplast, thylakoid–inner envelope contact sites (TIES) have been observed in TEM images of chloroplasts of land plants (Staehelin, 1986), algae (Gibbs, 1962), and in cyanobacteria (Mareš et al., 2019), and corroborated by modern cryo-electron tomography (Liang et al., 2018; Mai et al., 2019; Rast et al., 2019; Weiss et al., 2022; Wietrzynski et al., 2020). Biochemical studies show that lipid exchange between the inner envelope and thylakoids can occur in an ATP-independent and temperature-insensitive manner (Rawyler et al., 1992), further supporting a contact-based mechanism, where lipid transfer enzymes often function energy-independently (Lahiri et al., 2015). In vivo, thylakoids continue to form under conditions that inhibit vesicle fusion including low temperature (Krol et al., 1987) or mutation of *cpsfl1* (Hertle et al., 2020). These findings converge on a model in which vesicles and TIES operate in parallel to support thylakoid membrane biogenesis. We therefore hypothesized that TIES are structured by proteins essential for coordinating lipid flow to thylakoids.

To explore mechanisms coordinating lipid flow and membrane architecture within chloroplasts, we applied two parallel strategies: identification of chloroplast-targeted proteins with similarity to lipid transport or membrane structuring proteins, and proteomic profiling of intermediate-density chloroplast membranes. Together, these approaches revealed candidate proteins enriched at discrete chloroplast subdomains, with roles in shaping thylakoid-envelope organization.

## RESULTS

### Identification of Chloroplast Localized Lipid Transporters or Membrane Contact Site Homologs

To identify proteins potentially involved in lipid transport from the chloroplast inner envelope (IE) to the thylakoid membrane, we first conducted a domain homology-based screen targeting known MCS proteins and lipid transporters. In addition to sequence similarity, candidate genes were selected based on predicted chloroplast targeting or prior detection in chloroplast proteomic datasets. Preference was given to proteins with predicted lipid binding or membrane interaction. Ultimately, we refined our list to 19 tractable candidates for further investigation (Table S1).

To determine the subcellular locations of these candidate proteins, each was fused to monomeric Kusabira Orange kappa (mKOK, Figure 1A; Tsutsui et al., 2008), and transiently expressed in *Nicotiana benthamiana*. Proteins were visualized via confocal laser-scanning microscopy (Figure 1B). Two fusion constructs were used as positive controls: SENSITIVE TO FREEZING 2 (SFR2), a known outer envelope membrane protein (Roston et al., 2014) fused to enhanced green fluorescent protein, and CHLOROPLAST SEC14-LIKE 1 (CPSFL1), previously shown to localize to stromal vesicles (Hertle et al., 2020) fused to mKOK. Both controls exhibited strong spatial overlap with chlorophyll autofluorescence, consistent with chloroplast localization (Figure 1B). As a negative control for chloroplast localization, we used a fusion of the KAR2 signal peptide, enhanced cyan fluorescent protein, and the HDEL endoplasmic reticulum (ER) retention sequence (eCFP-HDEL; Cai et al., 2015). This construct showed minimal overlap with chlorophyll signal, indicating its non-chloroplast location. To quantitatively assess the degree of colocalization between fluorescent protein signals and chlorophyll autofluorescence, we used CellProfiler (Stirling et al., 2021b).

**Figure 1.**
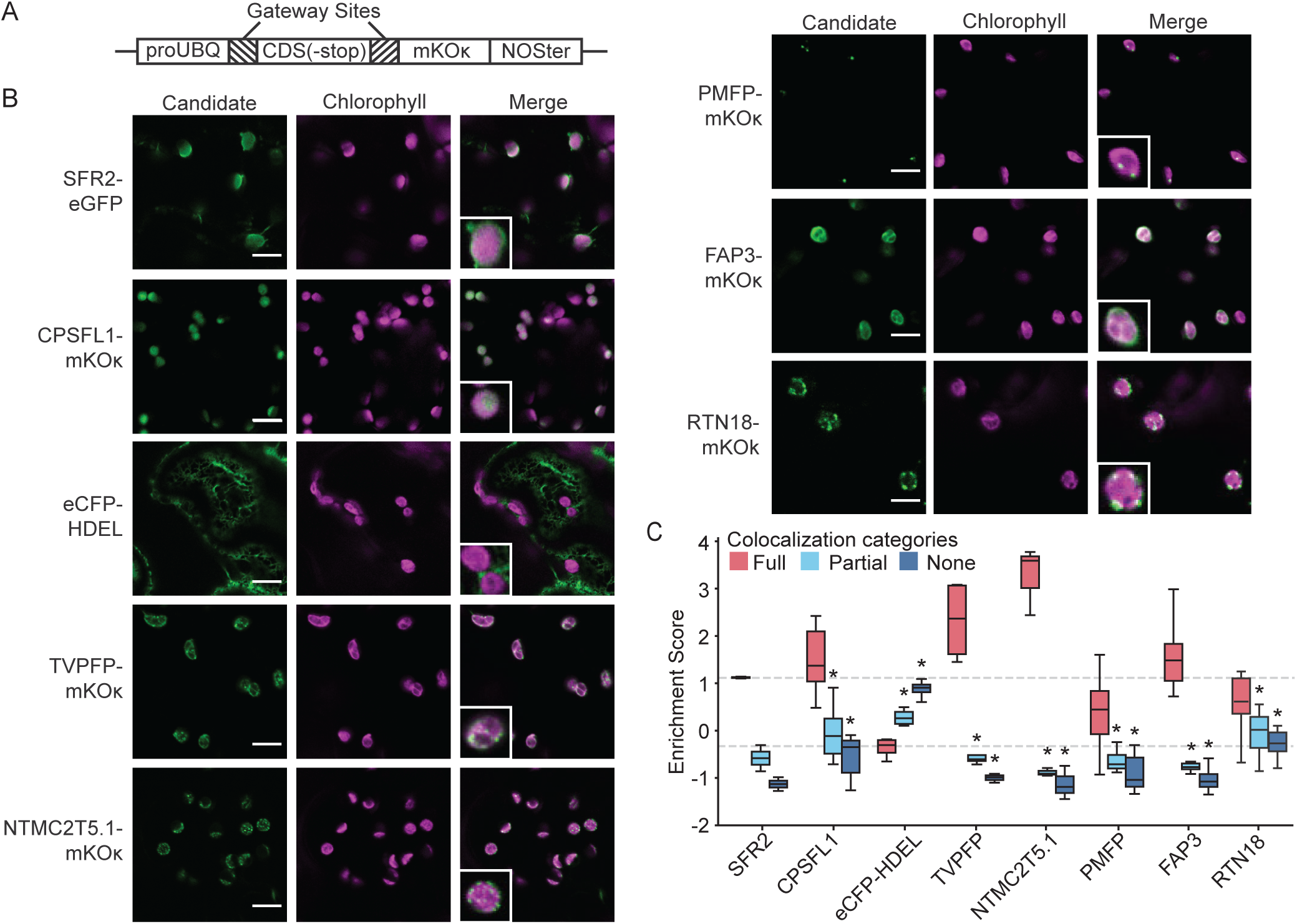
Chloroplast-localized candidate proteins colocalize with chlorophyll autofluorescence. (A) Expression constructs used for transient transformation of *N. benthamiana*. Candidate genes were C-terminally fused to mKOK and expressed under the UBQ10 promoter. (B) Confocal images showing fluorescence of candidate fusion proteins (green), chlorophyll autofluorescence (magenta), and merged channels. Positive controls (SFR2-eGFP, CPSFL1-mKOK) and a negative control (eCFP-HDEL) are included. Insets show zoomed 4 × 4 μm regions. Scale bar = 10 μm. (C) CellProfiler- based quantification of colocalization between candidate signals and chlorophyll autofluorescence. Colocalization categories are represented as boxplots (n 2: 3 trials, >100 chloroplasts total). Dashed lines indicate the median values for full colocalization of the positive (SFR2-eGFP) and negative (eCFP-HDEL) controls. Asterisks denote statistical significance (*p* < 0.05) by Welch’s ANOVA followed by Dunnett’s T3 multiple comparisons test between the full colocalization group and other categories for each protein (see Table S2 for full statistics). Abbreviations: ProUBQ, Ubiquitin 10 promoter; CDS, coding sequence; mKOK, monomeric Kusabira Orange kappa; NOSter, Nopaline synthase terminator; SFR2, SENSITIVE TO FREEZING 2; eGFP, enhanced green fluorescent protein; eCFP, enhanced cyan fluorescent protein; HDEL, ER retention signal; CPSFL1, Chloroplast Sec14-like 1; TVPFP, Tvp38 Family Protein; NTMC2T5.1, N-Terminal Transmembrane C2 Domain Type 5.1; PMFP, Plasma Membrane Fusion Protein; FAP3, Fatty Acid Binding Protein 3; RTN18, Reticulon 18.

Chloroplasts were categorized into three groups based on colocalization: full, partial, or no overlap (Figure 1C). As expected, the positive controls showed a high frequency of full colocalization, while the eCFP-HDEL negative control predominantly exhibited no colocalization (Figure 1C).

Five candidate proteins showed strong chloroplast colocalization: TVP38 FAMILY PROTEIN (TVPFP), N-TERMINAL TRANSMEMBRANE C2 DOMAIN TYPE 5.1 (NTMC2T5.1), PLASMA MEMBRANE FUSION PROTEIN (PMFP), and RETICULON 18 (RTN18), all displayed punctate fluorescence signals, while FATTY ACID BINDING PROTEIN 3 (FAP3) showed a diffuse signal pattern (Figure 1B, insets). These five candidates displayed a high frequency of chloroplasts fully colocalized with the fluorescent marker, comparable to the SFR2-eGFP positive control and significantly more than the frequency of partial or no colocalization (Figure 1C, Table S2).hese patterns suggest bona fide chloroplast localization rather than incidental spatial overlap. Additional confocal images of each candidate are provided in Figure S1.

Of the 19 candidate proteins tagged with mKOK, four showed subcellular localization patterns that were not primarily chloroplastic but were enriched near chloroplasts compared to controls (Figure 2A, Figure S2): PLECKSTRIN HOMOLOGY DOMAIN PROTEIN 1 (PHDP1), PLECKSTRIN HOMOLOGY DOMAIN PROTEIN 2 (PHDP2), CONSERVED OLIGOMERIC GOLGI COMPLEX 3 (COG3), and NON-SPECIFIC LIPID TRANSFER GPI-ANCHORED PROTEIN 20 (LTPG20). PHDP1 localized partially to chloroplasts but was also detected in other membrane compartments. PHDP2 and LPTG20 signals encircled chloroplasts but were similarly present in other cellular regions. COG3 had a diffuse signal that overlapped with chlorophyll signal frequently. CellProfiler quantification revealed that these proteins exhibited comparable frequencies across all colocalization classes, consistent with partial or heterogeneous chloroplast association (Figure 2B, Table S2).

**Figure 2.**
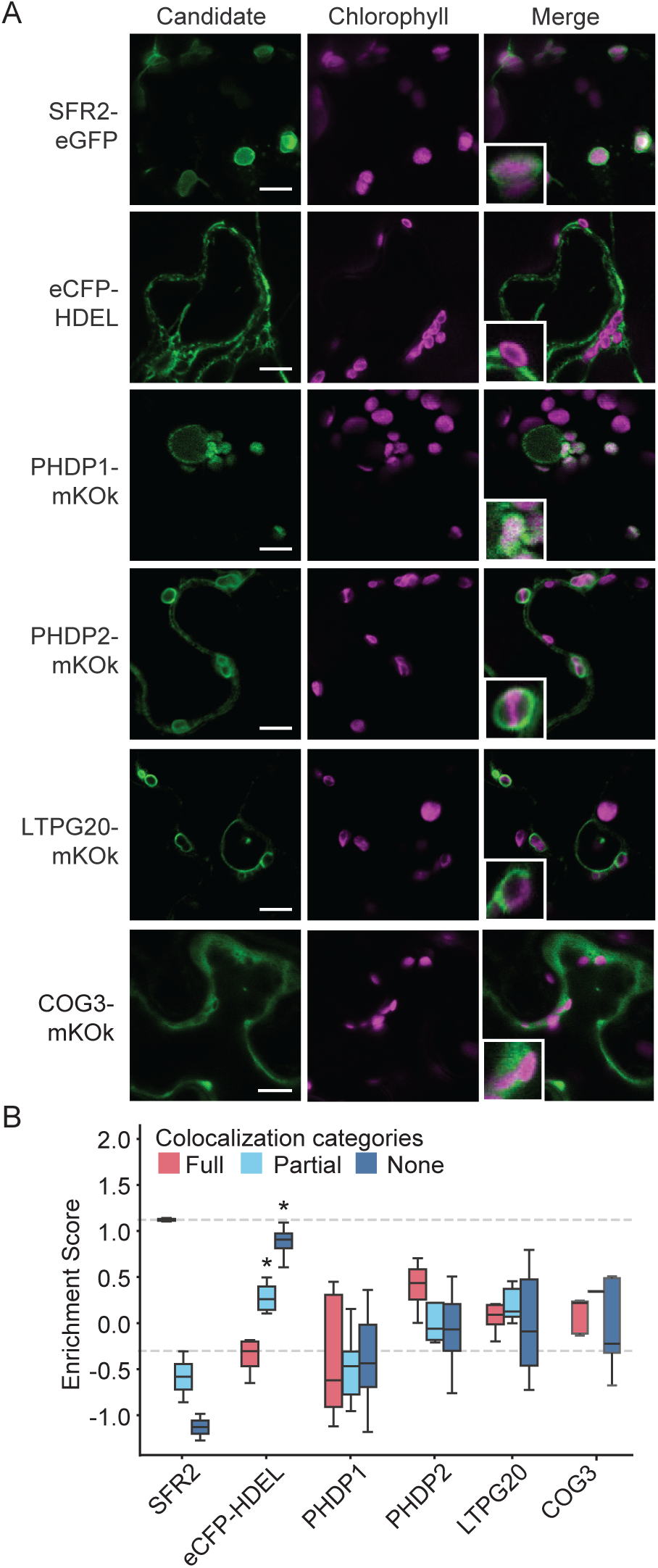
Candidate proteins with partial or peripheral chloroplast association. (A) Confocal images showing transient expression of candidate fusion protein fluorescence (green), chlorophyll autofluorescence (magenta), and merged channels. SFR2-eGFP and eCFP-HDEL serve as chloroplast-positive and -negative controls, respectively. Insets = 4 × 4 μm; scale bar = 10 μm. (B) CellProfiler-based quantification of colocalization between candidate signals and chlorophyll autofluorescence. Colocalization categories are represented as boxplots (n 2: 3 trials, >100 chloroplasts total). Dashed lines indicate the median values for full colocalization of the positive (SFR2-eGFP) and negative (eCFP-HDEL) controls. Asterisks denote statistical significance (*p* < 0.05) by Welch’s ANOVA followed by Dunnett’s T3 multiple comparisons test between the full colocalization group and other categories for each protein (see Table S2 for full statistics). Abbreviations: SFR2, SENSITIVE TO FREEZING 2; eGFP, enhanced green fluorescent protein; eCFP, enhanced cyan fluorescent protein; HDEL, ER retention signal; mKOK, monomeric Kusabira Orange kappa; PHDP1/2, Pleckstrin Homology Domain Protein; LTPG20, GLYCOSYLPHOSPHATIDYLINOSITOL-ANCHORED LIPID PROTEIN TRANSER 20; COG3, Conserved Oligomeric Golgi complex protein 3.

The remaining nine candidate proteins showed minimal overlap with chlorophyll autofluorescence and instead localized to other subcellular compartments (Figure S3A). All of these candidates had the highest frequency of no colocalization (Figure S3B, Table S2). Several, including CALCIUM-DEPENDENT LIPID-BINDING FAMILY PROTEIN (CALB) and OXYSTEROL BINDING PROTEIN-RELATED PROTEINS 3A, 3B, and 3C (ORP3A, ORP3B, ORP3C), sometimes surrounded chloroplasts or were enriched at regions near chloroplasts. These candidates showed increased partial colocalization (Figure S3B), suggesting possible peripheral association. The remaining candidates—DYNAMIN-RELATED PROTEIN 1A (DRP1A/ADL1), DYNAMIN-RELATED PROTEIN 3A (DRP3A/ADL2), RETICULON 17 (RTN17), SYNTAXIN OF PLANTS 21 (SYP21), and PUTATIVE LIPID BINDING PROTEIN (PLBP)—showed no detectable association with chloroplasts, presumably reflecting their localization to other subcellular compartments in this expression system.

### Isolation of Intermediate Density Chloroplast Membranes

Density-based membrane fractionation has been widely used to enrich for MCSs by isolating fractions with intermediate buoyant densities between associated membrane systems. This approach has been successfully applied to characterize contact sites between the mitochondrial inner and outer membranes (Hermann et al., 1998; Pon et al., 1989), mitochondria and the ER (Rusiñol et al., 1994; Vance, 2014), and bacterial inner and outer envelope membranes (Ishidate et al., 1986; Tefsen et al., 2005). To enrich for putative TIES in chloroplasts, we adapted a similar strategy using intact chloroplasts isolated from *Pisum sativum*.

Chloroplasts were hypertonically lysed and membranes were first separated by density using a discontinuous sucrose gradient. This resulted in crude outer envelope, inner envelope, and thylakoid membrane vesicle fractions. These fractions were then subjected to a second round of hypertonic lysis and homogenization to disrupt non-specific membrane associations and promote the isolation of membrane subdomains. The resulting membranes were resolved on a continuous sucrose gradient for finer separation (Figure 3A). To assess protein content and identify membrane-specific banding patterns, continuous gradient fractions from the second separation were analyzed by SDS-PAGE and visualized by silver staining (Figure 3B). Distinct banding patterns corresponding to IE and thylakoid membranes were consistently observed (asterisks, Figure 3B, compare lanes 11 & 12 in the upper panel with lanes 16 & 17 in the lower panel). Notably, fractions of intermediate density between the IE and thylakoid frequently exhibited unique banding patterns (Figure 3B, fractions 13 - 15), suggesting the presence of a distinct membrane population. This was most clearly observed in the light thylakoid fractions.

**Figure 3.**
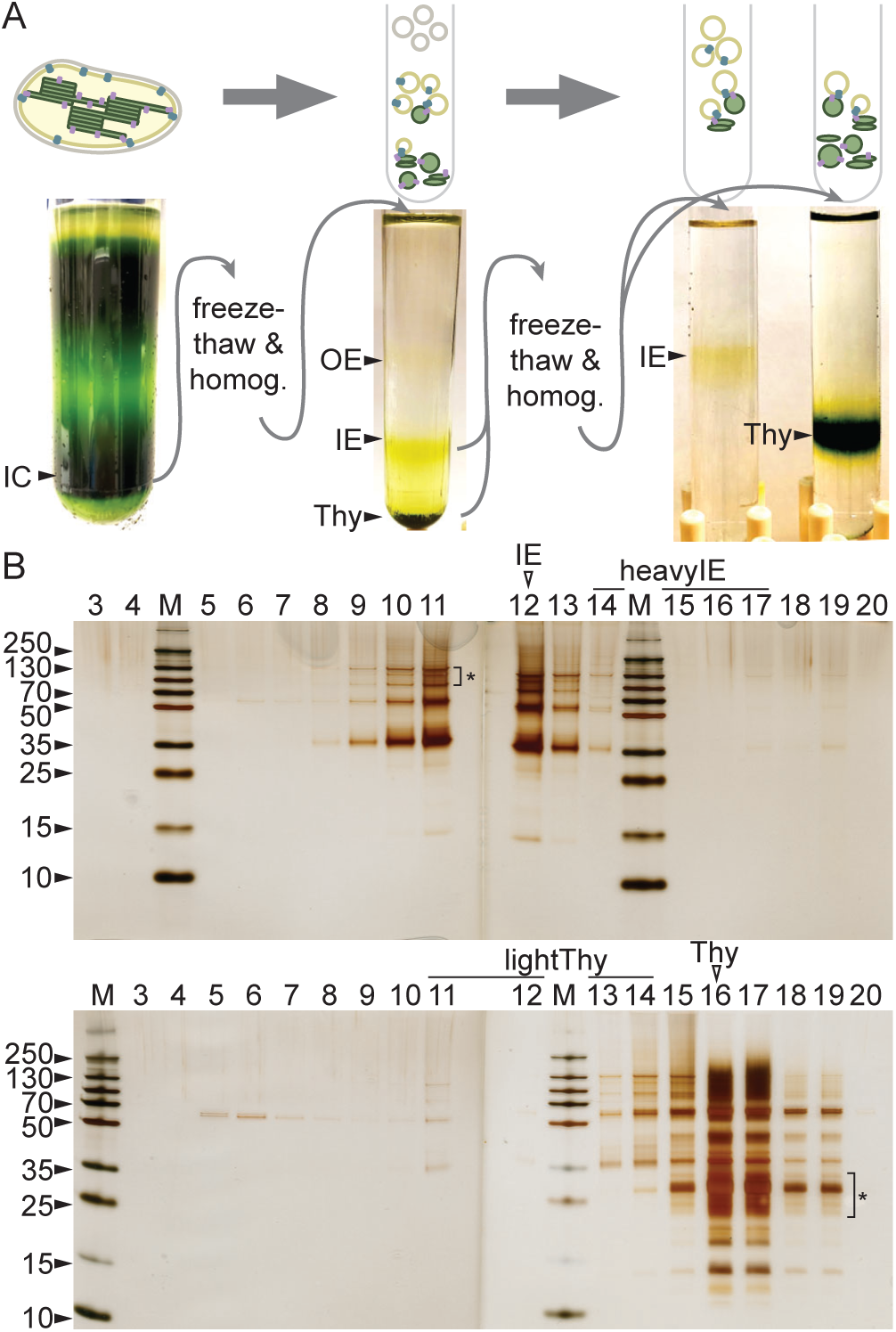
Fractionation of chloroplast membranes of intermediate density. (A) Workflow for chloroplast membrane fractionation. Intact chloroplasts (IC) were isolated, then homogenized into vesicles and fractionated by discontinuous sucrose gradients to enrich fractions containing outer envelope (OE), inner envelope (IE), and thylakoid (Thy) membranes. These fractions were homogenized again and finely resolved on continuous sucrose gradients to isolate inner envelope (IE), thylakoid (Thy), and intermediate-density fractions. (B) Silver-stained SDS-PAGE of continuous gradient fractions of the IE (top panel) or Thy (bottom panel) continuous gradients. Fraction numbers indicated at the top of the panel proceed in order from least-dense to most- dense of a continuous gradient. Asterisks highlight banding patterns that are characteristic of IE or Thy. Open arrowheads or black underline indicate fractions selected for proteomic analysis.

### Lipid Metabolic Enzymes Are Enriched in Light Thylakoid Fractions

To characterize the protein composition of intermediate-density chloroplast membrane fractions, we performed proteomic analysis on IE, heavy IE (heavyIE), thylakoid (Thy), and light thylakoid (lightThy) fractions. After analysis by LC-MS/MS, the final dataset included a total of 1,590 proteins confidently identified (with at least 2 spectral matches at 99% confidence, detected in 3 or more replicates of at least one fraction, and in 4 or more samples total, Table S3). Principal component analysis was applied to these samples to examine if the spread of proteomics data aligned with identities of unique membrane density fractions. The four membrane fractions were clustered with heavyIE and lightThy fractions positioned between well separated IE and Thy clusters (Figure 4A). This spatial separation suggests that heavyIE and lightThy represent discrete subpopulations of chloroplast membranes.

**Figure 4.**
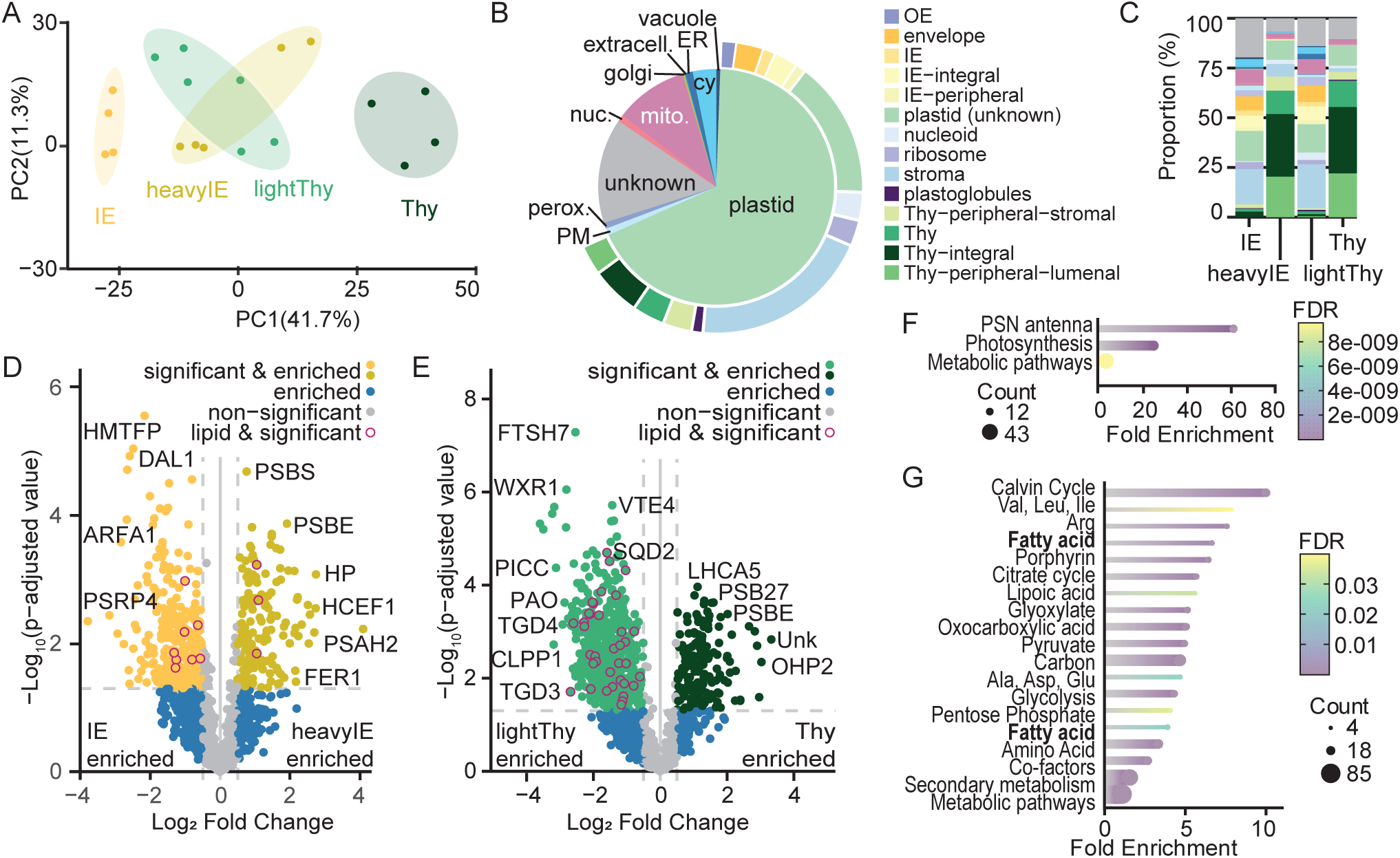
Proteomics reveals functional specialization of intermediate chloroplast membranes. (A) Principal component analysis (PCA) of proteomic profiles from four chloroplast membrane fractions shows clustering and separation of lightThy and heavyIE from canonical IE and thylakoid fractions. (B) Subcellular localization summary of all identified proteins (n = 1,590). (C) Patterns of plastid subcompartment-localized proteins enriched in fractions. Proteins were considered enriched if they exhibited a fold-change (|log21) 2: 0.5 and an adjusted p-value (–log10) 2: 1.15. (D–E) Volcano plots highlight proteins enriched in heavyIE (vs. IE) and lightThy (vs. Thy); a selection of significant proteins are labeled, all lipid metabolism proteins (AraLip database) are indicated. (F– G) KEGG pathway analysis of proteins enriched in heavyIE and lightThy. Dot size reflects number of proteins in the pathway; color indicates FDR. Abbreviations: KEGG, Kyoto Encyclopedia of Genes and Genomes; FDR, false discovery rate; cy, cytosol; ER, endoplasmic reticulum; extracell., extracellular; mito., mitochondria; nuc., nucleus; perox., peroxisome; PM, plasma membrane; PSRP4, PLASTID-SPECIFIC RIBOSOMAL PROTEIN 4; ARFA1, ADP-RIBOSYLATION FACTOR A1F; HMTFP, HEAVY METAL TRANSPORT/DETOXIFICATION SUPERFAMILY PROTEIN; DAL1, DIAP1-LIKE PROTEIN 1; PSBS, PHOTOSYSTEM II SUBUNIT S; PSBE, PHOTOSYSTEM II SUBUNIT E; HP, HYPOTHETICAL PROTEIN; HCEF1, HIGH CYCLIC ELECTRON FLOW 1; PSAH2, PHOTOSYSTEM I SUBUNIT H-2; FER1, FERRETIN 1; TGD3 OR 4, TRIGALACTOSYLDIACYLGLYCEROL 3 OR 4; CLPP1, CASEINOLYTIC PROTEASE PROTEIN 1; PAO, PHEOPHORBIDE A OXYGENASE; PICC, PAMP-INDUCED COILED COIL; WXR1, WEAK AUXIN RESPONSE1; FTSH7, FTSH PROTEASE 7; VTE4, GAMMA-TOCOPHEROL METHYLTRANSFERASE; SQD2, SULFOQUINOVOSYLDIACYLGLYCEROL 2; LHCA5, PHOTOSYSTEM I LIGHT HARVESTING COMPLEX GENE 5; PSB27, PHOTOSYSTEM II 11 KDA PROTEIN- LIKE PROTEIN; UNK, UNKNOWN; OHP2, ONE-HELIX PROTEIN 2.

To evaluate the quality of the dataset, we next looked at the subcellular localization of identified proteins using manually curated annotations from PPDB (Sun et al., 2009) and SUBA (Hooper et al., 2022). Of the 1,590 proteins identified, 1,560 had Arabidopsis homologs and could be investigated. Of these, 1,161 (74.4%) were classified as plastid-localized (Figure 4B, Table S4), suggesting successful chloroplast isolation and fractionation. Independent assessment using SUBA’s Multiple Marker Abundance Profiling tool (Hooper et al., 2022) estimated a 90.7% plastid enrichment based on relative numbers of high-confidence markers, in agreement with our manual analysis.

We then assessed the composition of each membrane fraction by examining the distribution of proteins assigned to specific plastid subcompartments. Proteins significantly enriched in each fraction were identified using a log2 fold change threshold of ±0.5 and an adjusted p-value cutoff of –log10(p) 2: 1.15. These criteria yielded 224 proteins enriched in the IE fraction, 143 in heavyIE, 491 in lightThy, and 159 in the thylakoid fraction (Table S4). Suborganellar localizations for plastid proteins were then manually curated using annotations from PPDB, SUBA, and published literature. While the suborganellar localization of some proteins remains unresolved, sufficient annotation was available to discern distinct enrichment patterns across the fractions (Figure 4C). As expected, the IE fraction was enriched in envelope proteins (21%) relative to the thylakoid fraction (1%), while the thylakoid fraction was enriched in thylakoid proteins (73%) compared to the IE fraction (6%). The lightThy fraction was enriched for envelope (24%), unclassified plastid (14%), and non-plastid proteins (30%), and contained only 4% thylakoid proteins. This profile stands in contrast to the thylakoid fraction, which contained 1% envelope, 10% unclassified plastid, 13% non-plastid, and 73% thylakoid proteins. Similarly, the heavyIE fraction exhibited a complementary pattern to the canonical IE: it was enriched in thylakoid proteins (71%) but depleted in envelope (7%) and non-plastid proteins (10%) compared to the IE fraction (6% thylakoid, 21% envelope, 36% non-plastid, respectively). Together, these data demonstrate that lightThy and heavyIE are compositionally distinct from canonical IE and thylakoid membranes, with unique combinations of protein localizations.

To evaluate enrichment of MCS components, we compared Arabidopsis gene homologs from each fraction to the MCS database (Pan et al., 2024). The lightThy was notably enriched in proteins with homology to known MCS components: 128 proteins (79%) uniquely enriched in lightThy had significant similarity (Table S5). In comparison, fewer MCS homologs were identified in the IE (35%), heavyIE (16%), and Thy (13%) fractions.

These results suggest that the lightThy fraction was enriched in proteins associated with MCS.

To assess distribution of lipid metabolism, we mapped Arabidopsis gene identifiers to the Acyl-Lipid Metabolism database (Li-Beisson et al., 2013). The lightThy fraction again stood out, containing 30 lipid-related enzymes—more than three times as many as any other fraction (heavyIE: 3, IE: 9; Table 1). This enrichment was also evident in volcano plots indicating lipid metabolic proteins (Figures 4D and 4E). In contrast, the Thy fraction lacked detectable lipid metabolic enzymes, consistent with longstanding observations that lipid biosynthesis does not occur in thylakoid membranes (Douce, 1974; Seifert and Heinz, 1992).

**Table 1.** Lipid Metabolism Proteins Enriched in Chloroplast Membrane Fractions. This table lists lipid metabolism proteins identified by proteomic analysis as enriched in specific chloroplast membrane fractions. AGI refers to the Arabidopsis Gene Identifier corresponding to the closest BLAST match for the identified pea protein. Enriched indicates the membrane fraction(s) where the protein was significantly enriched. Name provides a brief functional annotation based on TAIR or recent literature. Abbrev. gives an abbreviation or commonly used name for the protein. References cite supporting studies demonstrating each protein’s role in lipid metabolism, drawn from the Acyl-Lipid Metabolism database (Li-Beisson et al., 2013).

| AGI | Enriched | Name | Abbrev. | Reference(s) |
| --- | --- | --- | --- | --- |
| AT1G77590 | IE | Long-Chain Acyl-CoA Synthetase | LACS9 | (Shockey et al., 2002) |
| AT2G01320 | IE, lightThy | ABC Transporter | WBC7 / ABCG7 |  |
| AT2G30200 | IE | Malonyl-CoA : ACP Malonyltransferase | MCMT | (Bryant et al., 2011; Zhang et al., 2003) |
| AT2G38040 | IE, lightThy | Carboxyltransferase alpha Subunit of Heteromeric ACCase | ACCDα | (Bilder et al., 2006; Reverdatto et al., 1999) |
| AT2G38670 | IE | CTP:phosphorylethanolamine cytidyltransferase | PECT1 | (Mizoi et al., 2006) |
| AT3G05510 | IE, lightThy | Cardiolipin Transacylase | TAZ / TAZ2 | (Y. Xu et al., 2003) |
| AT3G11170 | IE, lightThy | Linoleate Desaturase | FAD7 | (Browse et al., 1986; Iba et al., 1993; Matsuda et al., 2005) |
| AT3G25780 | IE | Allene Oxide Cyclase | AOC3 |  |
| ATCG00500 | IE, lightThy | Carboxyltransferase beta Subunit of Heteromeric ACCase | ACCDβ | (S. J. Li et al., 1992) |
| AT1G01090 | heavyIE, lightThy | E1a component of Pyruvate Dehydrogenase Complex | PDH (E1 alpha) | (De Meirleir et al., 1992; Johnston et al., 1997) |
| AT2G34590 | heavyIE, lightThy | E1b component of Pyruvate Dehydrogenase Complex | PDH (E1 beta) | (R. M. Brown et al., 2004; Johnston et al., 1997) |
| AT3G16950 | heavyIE, lightThy | Dihydrolipoamide Dehydrogenase, E3 component of Pyruvate Dehydrogenase Complex | LPD1 (E3) | (Chen et al., 2010; Engels & Pistorius, 1997) |
| AT1G08640 | lightThy | Chloroplast J-like Domain 1 | CJD1 | (Ajjawi et al., 2011) |
| AT1G19800 | lightThy | Permease-like Protein of Inner Chloroplast Envelope | TGD1 | (C. Xu et al., 2003) |
| AT1G34430 | lightThy | E2 component of Pyruvate Dehydrogenase Complex | EMB300 3 (E2) | (Lin et al., 2003) |
| AT1G65260 | lightThy | Vesicle-Inducing Protein in Plastids | VIPP1 | (Aseeva et al., 2004; Kroll et al., 2001; Westphal et al., 2001) |
| AT1G65410 | lightThy | ATPase | TGD3 | (Lu et al., 2007) |
| AT1G70670 | lightThy | Caleosin | CLO4 | (Naested et al., 2000) |
| AT1G70680 | lightThy | Caleosin | CLO6 | (Naested et al., 2000) |
| AT1G74960 | lightThy | Ketoacyl-ACP Synthase II | KASII | (Carlsson et al., 2002; Huang et al., 1998; Pidkowich et al., 2007) |
| AT1G77590 | lightThy | Long-Chain Acyl-CoA Synthetase | LACS9 | (Shockey et al., 2002) |
| AT2G05990 | lightThy | Enoyl-ACP Reductase | ENR1 (MOD1) | (Mou et al., 2000; Rafferty et al., 1995) |
| AT2G11810 | lightThy | Monogalactosyldiacylglycerol Synthase | MGD3 | (Awai et al., 2001; Benning & Ohta, 2005; Kobayashi et al., 2009) |
| AT2G39290 | lightThy | Phosphatidylglycerol-Phosphate Synthase | PGP1 | (Babiyshuk et al., 2003; Hagio et al., 2002; Müller & Frentzen, 2001; C. Xu et al., 2002) |
| AT3G06510 | lightThy | galactolipid:galactolipid galactosyltransferase | SFR2 | (Moellering et al., 2010; Thorlby et al., 2004) |
| AT3G06960 | lightThy | Protein Involved in Lipid Transport | TGD4 | (C. Xu et al., 2008) |
| AT3G10840 | lightThy | Lysophospholipase | n.a. |  |
| AT3G20320 | lightThy | Phosphatidic Acid-Binding Protein | TGD2 | (Awai et al., 2006) |
| AT3G25860 | lightThy | Dihydrolipoamide Acetyltransferase, E2 component of Pyruvate Dehydrogenase Complex | LTA2 (E2) | (Lin et al., 2003; Mooney et al., 1999) |
| AT4G02510 | lightThy | plastid protein import 2 | TOC159 | (Afithile et al., 2013) |
| AT5G01220 | lightThy | UDP-sulfoquinovose:DAG sulfoquinovosyltransferase | SQD2 | (Yu et al., 2002) |
| AT5G05580 | lightThy | Linoleate Desaturase | FAD8 | (Matsuda et al., 2005; McConn et al., 1994) |
| AT5G10160 | lightThy | Hydroxyacyl-ACP Dehydrase | HAD | (A. Brown et al., 2009) |
| AT5G16390 | lightThy | Biotin Carboxyl Carrier Protein of Heteromeric ACCase | BCCP1 | (Choi et al., 1995; X. Li et al., 2011; Reverdatto et al., 1999) |

To further investigate functional specialization, we performed KEGG pathway and GO enrichment analysis on the fractions. The heavyIE fraction was enriched for photosynthesis-related pathways, including those involving the photosystem antenna complex (Figure 4F, Table S6). In contrast, the lightThy fraction showed broader enrichment in both hydrophobic and hydrophilic metabolic processes, including fatty acid, porphyrin, and lipoic acid metabolism, as well as portions of the glycolysis pathway and the Calvin–Benson–Bassham cycle (Figure 4G, Table S6).

### Tagging and Transient Expression of Candidate Genes Identified by Proteomics

To confirm the chloroplast distribution of proteins found to be enriched in the lightThy fractions, three additional proteins were selected for fluorescent tagging with mKOK and transient expression in *Nicotiana benthamiana*. In addition to their significant enrichment in the lightThy fraction, these candidates were selected based on prior functional evidence linking them to lipid trafficking or MCS. TRIGALACTOSYLDIACYLGLYCEROL 3 (TGD3) is a component of the inner envelope-localized TGD lipid transport complex (Roston et al., 2012) important for lipid trafficking between chloroplast envelope membranes (Lu et al., 2007). TGD3 was enriched in both heavyIE and lightThy fractions. LETM1-LIKE (LETM1L) is a putative membrane protein homologous to the mitochondrial LetM1, which regulates cristae morphology and contact site formation in animal cells (Nakamura et al., 2020). VESICLE-INDUCING PROTEIN IN PLASTIDS 1 (VIPP1) is a widely conserved chloroplast protein homologous to bacterial PspA, and has been observed near thylakoid biogenesis sites (Gupta et al., 2021). All three proteins showed strong colocalization with chlorophyll autofluorescence when expressed in tobacco leaves (Figure 5A). Additionally, signal from each protein exhibited non-uniform distribution relative to chlorophyll fluorescence, with TGD3 being the most diffuse, while VIPP1 and LETM1L formed clear punctate substructures (Figure 5A, insets). Together, these results validate that the lightThy fraction is enriched with chloroplast proteins distributed at discrete locations within the chloroplast (Figure 5B) and distinct from typical envelope localization patterns (Figure 1B) or chlorophyll fluorescence of thylakoid membranes.

**Figure 5.**
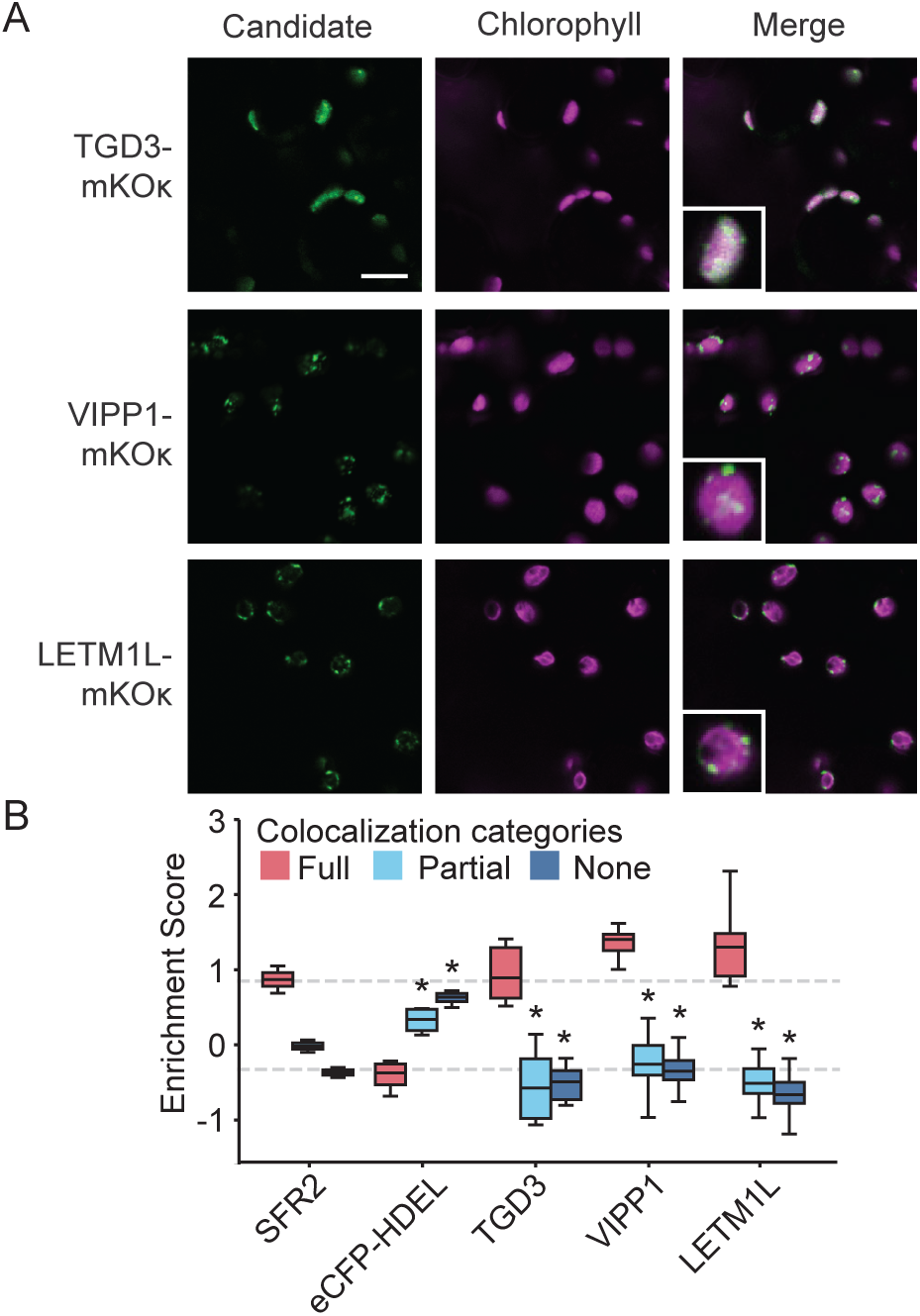
Candidate proteins from intermediate density chloroplast fractions localize to discrete regions. (A) Confocal microscopy images of tagged candidates TGD3, VIPP1, and LETM1L showing fluorescence (green), chlorophyll signal (magenta), and merged images. Insets = 4 × 4 μm; scale bar = 10 μm. (B) CellProfiler-based quantification of colocalization between candidate signals and chlorophyll autofluorescence. Colocalization categories are represented as boxplots (n 2: 3 trials, >100 chloroplasts total). Dashed lines indicate the median values for full colocalization of the positive (SFR2-eGFP) and negative (eCFP-HDEL) controls. Asterisks denote statistical significance (*p* < 0.05) by Welch’s ANOVA followed by Dunnett’s T3 multiple comparisons test between the full colocalization group and other categories for each protein (see Table S2 for full statistics). Abbreviations: TGD3, TRIGALACTOSYLDIACYLGLYCEROL 3; VIPP1, VESICLE INDUCING PROTEIN IN PLASTIDS 1; LETM1L, LETM1-LIKE; mKOK, monomeric Kusabira Orange kappa.

### Loss of TVPFP and PMFP Affects Chloroplast Membrane Ultrastructure

Two candidates identified in our initial confocal microscopy screen, TVPFP and PMFP, are previously uncharacterized proteins that exhibited punctate fluorescent signals overlapping with chlorophyll autofluorescence (Figure 1B), reminiscent of VIPP1 and LETM1L (Figure 5A). They were selected for additional characterization based on their predicted roles and unstudied nature. TVPFP is a member of the evolutionarily conserved Tvp38/DedA domain-containing family of transmembrane proteins, which in yeast and mammals have been identified as lipid scramblases assisting in lipid transport at ER–autophagosome contact sites (Ghanbarpour et al., 2021; Huang et al., 2021; Li et al., 2021). PMFP belongs to the plasma membrane fusion protein family (PTHR36398), a group of proteins conserved throughout the green lineage but lacking functional characterization (Mi et al., 2021). TVPFP is predicted to be a multi-pass membrane protein, while PMFP contains two predicted monotopic helices (Figure 6A).

**Figure 6.**
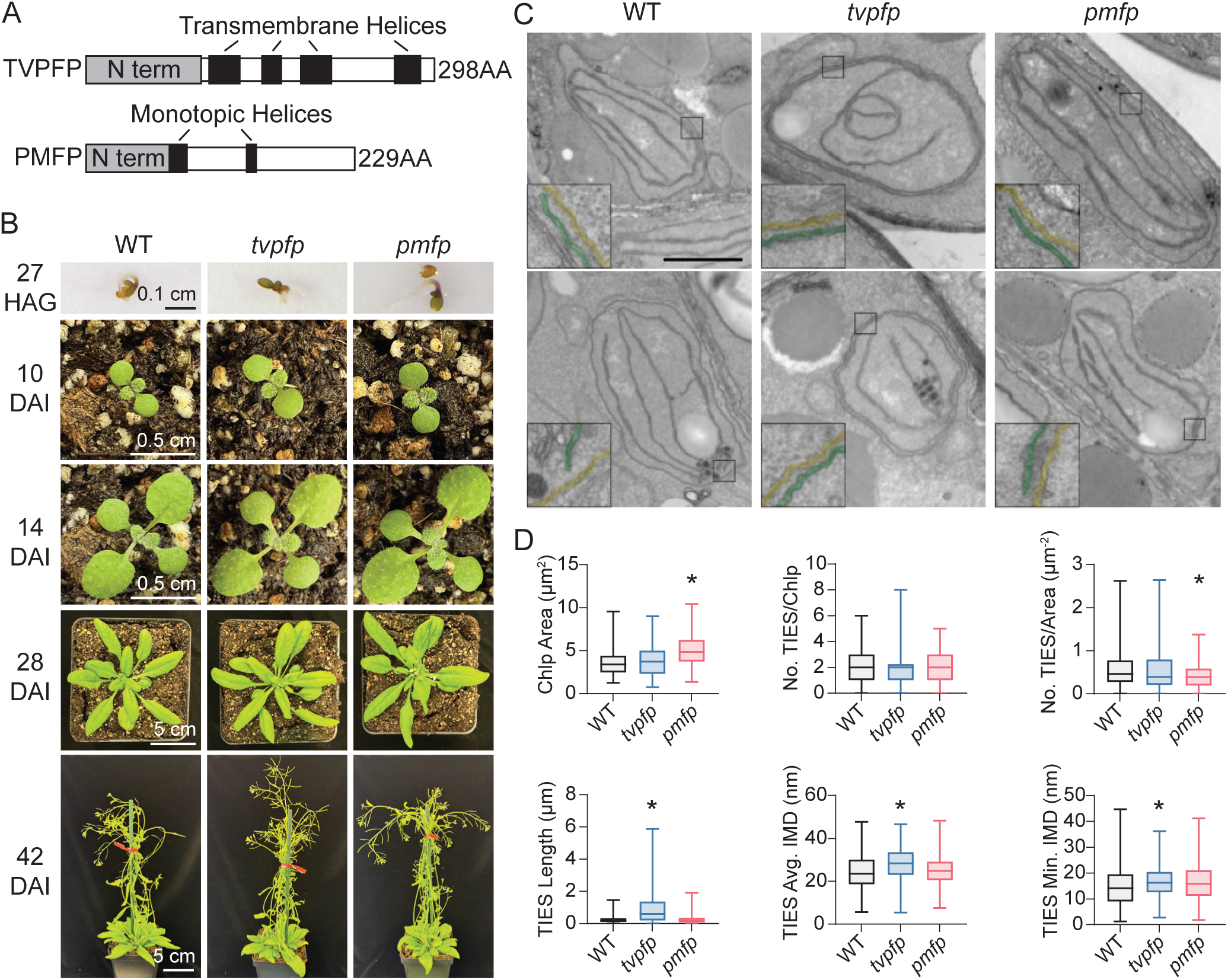
Mutants lacking TVPFP or PMFP show altered chloroplast membrane architecture. (A) AlphaFold-predicted membrane models of TVPFP and PMFP show structural features supporting membrane integration or association. (B) Growth phenotypes of wild-type and mutant plants under standard conditions. HAG, hours after germination; DAI, days after imbibition. (C) Transmission electron micrographs of chloroplast ultrastructure in cotyledons of seedlings germinated with continuous light (scale = 1 μm). Inset (4x zoom) shows area of closely apposed thylakoid (green) to inner envelope (yellow). (D) Quantification of chloroplast area, number of TIES per chloroplast and per area, TIES length, and intermembrane distances (IMD). Asterisks indicate significance (p :5 0.05); >100 chloroplasts per genotype analyzed.

To test whether the presence of TVPFP and PMFP affected the association of the chloroplast IE and thylakoid membranes, we investigated T-DNA insertion loss-of-function lines for each in *Arabidopsis thaliana.* Homozygous mutants for both genes were confirmed by PCR (Figure S4). Under standard growth conditions, both *tvpfp* and *pmfp* plants developed normally and reached reproductive maturity, indicating the absence of strong growth defects (Figure 6B). To examine the role of these genes in chloroplast membrane organization, we germinated seedlings under continuous light—a condition that induces direct proplastid-to-chloroplast differentiation and stimulates lipid-dependent thylakoid expansion (Liang et al., 2018). Cotyledon plastid development was assessed by TEM (Figure 6C, Figure S5). Intriguingly, in *tvpfp* mutants, thylakoids were frequently observed to associate with the envelope membranes for most of the section. Quantification of multiple TIES features confirmed that in *tvpfp*, there was a significant increase in TIES length of more than 3 times that of WT (WT, 0.28 μm; *tvpfp,* 1.02 μm; Figure 6D, Table S7). Mutants of *pmfp* were reduced by about 25% compared to WT in the number of TIES per chloroplast area (WT, 0.57 μm^-2^; *pmfp,* 0.42 μm^-2^; Figure 6D, Table S7). Together, these results are consistent with the involvement of both TVPFP and PMFP in affecting chloroplast membrane architecture at thylakoid-inner envelope interfaces.

## DISCUSSION

In this study, we identified chloroplast-localized proteins with potential roles in thylakoid membrane biogenesis using two complementary strategies: homology-based candidate selection and proteomic profiling of fractionated chloroplast membranes. We identified several proteins—including TVPFP, PMFP, RTN18, and NTMC2T5.1—localized to discrete puncta within chloroplasts, consistent with their involvement in specialized suborganellar functions (Figure 1). Proteomic profiling of chloroplast membranes intermediate in density between thylakoid and inner envelope revealed a distinct fraction enriched in lipid metabolic enzymes and transporters (Figure 4, Table 2), consistent with our hypothesis that this fraction represents a relevant zone for lipid exchange. We further tested functional roles of TVPFP and PMFP, showing that their absence in Arabidopsis resulted in perturbed chloroplast membrane ultrastructure (Figure 6). These results highlight the functional importance of previously uncharacterized proteins in shaping chloroplast membrane architecture and support the existence of specialized regions for lipid exchange.

Chloroplast envelope and thylakoid membranes are notoriously difficult to isolate without cross-contamination, as shown by the persistent detection of chlorophyll precursors in envelope preparations (Pineau et al., 1993), lipid biosynthetic activity in thylakoids (Dorne et al., 1990; Douce, 1974), and shared protein identities across suborganellar proteomes (Bouchnak et al., 2019; Ferro et al., 2003). We leveraged this overlap to isolate intermediate-density fractions—most notably, a “light thylakoid” fraction (Figure 4C, E, G) which was enriched in lipid transporters and biosynthetic enzymes (Table 1) and homologs of membrane contact site proteins (Table S5). Unlike earlier methods of digitonin-based thylakoid grana subfractionation (Tomizioli et al., 2014) or plastoglobule flotation (Shivaiah et al., 2022), our two-step homogenization and density centrifugation approach prioritized tightly membrane-associated proteins. Unlike the lightThy fraction, the heavyIE fraction was skewed toward photosynthetic machinery (Figure 4C, D, F). This asymmetric enrichment may reflect the relative abundance of thylakoids versus IE membranes, enabling a more refined physical separation of functional subdomains.

Strikingly, the lightThy fraction was enriched in multiple lipid transporters, lipid, chlorophyll, and isoprenoid biosynthetic pathways. Four components of the TRIGALACTOSYLDIACYLGLYCEROL (TGD) complex—TGD1 through TGD4—were present, consistent with this complex’s established role in lipid transfer between the ER and the chloroplast envelope (Fan et al., 2015). Similarly, FATTY ACID EXPORT 2 (FAX2), a member of the chloroplast envelope-localized fatty acid transporter family (Li et al., 2020; Tian et al., 2019), was also enriched. In addition, enzymes such as GERANYLGERANYL REDUCTASE (GGDR) and CHLOROPHYLL SYNTHETASE (CHLG)—both previously reported in thylakoid and envelope proteomes (Ferro et al., 2010; Peltier et al., 2004; Soll et al., 1983))—were enriched in the heavyIE fraction. Together, these findings support the idea that the lightThy fraction represents a specialized membrane interface enriched in hydrophobic metabolism and transport functions.

Several of the candidate proteins we investigated display features consistent with roles in chloroplast membrane architecture and lipid transport. TVPFP is a member of the evolutionarily conserved DedA/Tvp38 family, whose homologs in bacteria and mammals act as lipid scramblases and contact site regulators (Boughner and Doerrler, 2012; Huang et al., 2021). In Arabidopsis, *tvpfp* mutants exhibited significantly longer thylakoid-inner envelope contacts and increased intermembrane spacing (Figure 6D), phenotypes that mirror those observed in other systems when contact site tethering is dysregulated (Tábara and Escalante, 2016; Zhao et al., 2017). PMFP, a plant-conserved but uncharacterized protein, was detected in thylakoid proteomes (Ferro et al., 2010; Trotta et al., 2019) and localized to chloroplasts in our assays (Figure 1B). Its loss led to an increase in chloroplast area and a modest reduction in contact site density (Figure 6D). Though less severe than *tvpfp*, these changes indicate functional relevance in membrane organization. LETM1-LIKE, a homolog of mitochondrial contact site protein LETM1 (Nakamura et al., 2020), and VIPP1, previously implicated in chloroplast vesicle formation and membrane remodeling (Junglas et al., 2025; Kroll et al., 2001), were also chloroplast-localized and enriched in MCS-like fractions. These candidates, each localized to discrete chloroplast puncta and enriched in a fraction dense with lipid-related and MCS-homologous proteins, strengthen the case for a structured and functionally specialized intermembrane zone within chloroplasts.

Protein presence at punctate chloroplast subdomains (Figures 1,5) and the enrichment of lipid enzymes (Figure 4D, E, Table S4), specialized chemistry (Figure 4G, Table S6), and contact site homologs (Table S5) in the lightThy fraction is consistent with recent publications in Chlamydomonas (Sun et al., 2025; Wang et al., 2023) that identify functionally specialized regions of algal chloroplasts. These findings advance our understanding of land plant chloroplast subcompartmentalization and highlight new molecular tools for dissecting membrane dynamics critical for photosynthetic function.

## MATERIALS & METHODS

### Plant Materials and Growth Conditions

*Arabidopsis thaliana* ecotype Columbia was used as the wild-type background. Homozygous T-DNA insertion lines *tvpfp* (SALK_095443.29.45.x) and *pmfp* (SALK_149886.49.50.x) were obtained from the Arabidopsis Biological Resource Center (Alonso et al., 2003). Seeds were sown on soil, or sterilized and sown on sterile media, and stratified as described previously (Shomo et al., 2024). Plants were grown in a 22°C/18°C, 18 h/6 h light/dark cycle at 120 µmol m^-2^s^-1^ light intensity. *Pisum sativum* (’Little Marvel’) seedlings were grown for 2 weeks in vermiculite under similar light and temperature conditions prior to chloroplast isolation. *Nicotiana benthamiana* was grown with similar light and temperature conditions on Berger BM2 germination mix for approximately 4 weeks prior to infiltration.

### Cloning and Transient Expression

Candidate open reading frames were amplified from cDNA synthesized using oligo(dT) primers and total RNA from 4-week-old *A. thaliana*. PCR was performed using Q5 DNA polymerase (NEB) with gene-specific primers adding attB sites and omitting stop codons. PCR products were gel-purified and cloned into pDONR221, then recombined into pUBC(k)-DEST-mKOK using Gateway cloning (Invitrogen). Final constructs were sequence verified and transformed into *Agrobacterium tumefaciens* GV3101(pMP90) via electroporation (Lin, 1995). Verifed pUBC(k)-candidate:mKOk clones are available from Addgene. Cultures were grown in YEBS with antibiotics (Davis et al., 2009) and co-infiltrated into *N. benthamiana* leaves alongside pCB301-p19 to enhance expression (Win & Kamoun, 2004). Infiltration was essentially as described (Norkunas et al., 2018) with minor modifications: Agrobacteria were resuspended to OD600 = 0.5 in infiltration buffer (10 mM MES, pH 5.6, 10 mM MgCl2, 200 μM acetosyringone, 0.005% Silwet L-77) before infiltration. A strain carrying a fluorescent construct was combined 1:1 with the strain containing pCB301-p19 and incubated for 2 hours at room temperature with slight agitation. Third and fourth upper leaves of 4-week-old *N. benthamiana* were infiltrated using a needle-less syringe through small holes pricked in the leaf surface. Plants were incubated under 160 µmol m^-2^s^-1^ light for 2 h, dark for 24 h, then returned to growth conditions.

### Confocal Microscopy and Image Analysis

Leaf samples were imaged 2 days post-infiltration using a Nikon NiE-A1 confocal microscope with a 60× water immersion objective. Excitation/emission settings were as follows: eCFP (445/464–499 nm), eGFP (488/500–550 nm), mKOK (561/570–620 nm), and chlorophyll autofluorescence (640/662–737 nm). Images were captured using 2× line averaging. Single Z-plane confocal images were screened manually for frames with multiple in-focus chloroplasts. Images were pre-processed in ImageJ (Schneider et al., 2012) to adjust contrast, reduce noise, and convert to .tiff format. Pre-processed images were analyzed in CellProfiler (v4.2.7) (Stirling et al., 2021b) using a custom pipeline that segmented chloroplast signal via Otsu thresholding and quantified intensity across channels. A machine learning model was trained in CellProfiler Analyst (v3.0.4) (Stirling et al., 2021a) to classify images into three colocalization categories: full, partial, and none, based on fluorescence overlap between chlorophyll and candidate markers. Fast Gentle Boosting and Random Forest classifiers were used to optimize feature selection and reduce false positives. The final model had a classification accuracy of 94.8% based on confusion matrix evaluation. Output hit tables (.csv) were analyzed in Python (v3.12.6) using seaborn (v0.13.2; Waskom, 2021), pandas (McKinney, 2011), and matplotlib (v 3.8.4; Hunter, 2007). Statistical comparisons were performed using Welch’s ANOVA with Dunnet’s T3 post-hoc testing against positive (SFR2) and negative (eCFP-HDEL) controls.

### Chloroplast Isolation and Membrane Fractionation

Chloroplasts were isolated from *P. sativum* as described (Bruce et al., 1994), with all buffers supplemented with protease inhibitors (1 mM PMSF, 0.5 mM £-aminocaproic acid, and 1 mM benzamidine). Intact chloroplasts were washed and lysed by freeze-thaw and Dounce homogenization of 30 strokes using a tight pestle. Membranes were separated on a discontinuous sucrose gradient (0.46 M, 0.8 M, 1.0 M sucrose in TE buffer), followed by ultracentrifugation (180,000 × g, 3 h) as in Keegstra and Yousif (1986). IE and thylakoid fractions were recovered, diluted, and pelleted (130,000 × g, 30 min). Each was then adjusted to 0.2 M sucrose, and homogenization by freeze-thaw and Dounce was repeated before loading on a continuous sucrose gradient (0.73–1.8 M) formed by passive diffusion (Stone, 1974) and spun at 113,000 × g for 15 h. Fractions of 500 μL each were serially collected for analysis.

### Protein Extraction and Immunoblotting

Proteins were separated using Bolt 12% Bis-Tris gels and transferred to PVDF membranes. Membranes were blocked in EveryBlot buffer (BIO-RAD) and probed with primary antibodies against GFP (53061S, Cell Signaling), Tic110, PC, and OE33 (AD08 293, AS06 141, A806 142, Agrisera).

Secondary detection used HRP-conjugated anti-rabbit IgG (VL313544, Invitrogen) and Clarity Max ECL substrate (Bio-Rad), imaged on an Odyssey Fc system (LI-COR).

### Silver Staining and Gel-Based Protein Preparation

Gels were silver-stained using Pierce™ Silver Stain for Mass Spectrometry (Thermo), following the manufacturer’s instructions. Four biological replicates with high-quality membrane fractionation were processed in parallel. Fractions included the highest protein abundance IE and thylakoid fraction, as well as heavyIE and lightThy fractions as indicated (Figure 3B). As needed, heavy or light fractions were grouped to allow analysis of equal total proteins from each fraction. Proteins were first precipitated with trichloroacetic acid, normalized by Bradford assay, and then loaded into the top portion of Bolt 12% Bis-Tris gels. For in-gel digestion, protein bands were excised, washed, reduced, alkylated, and digested overnight with trypsin. Peptides were extracted, dried, and resuspended in 5% acetonitrile, 0.05% trifluoroacetic acid.

### LC–MS/MS and Data Analysis

Peptides were analyzed by nano-LC-MS/MS using a Waters CSH C18 column online to a Thermo Orbitrap Eclipse mass spectrometer. Spectra were searched using Mascot Server (v2.7.0, www.matrixscience.com) with a fragment ion mass tolerance of 0.060 Da and a parent ion tolerance of 15.0 ppm. Variable modifications included deamidation of asparagine and glutamine, oxidation of methionine, and carbamidomethylation of cysteine. Searches were performed against the *Pisum sativum* v1a annotated protein database (57,835 entries) and a common contaminants database cRAP_20150130.fasta (www.thegpm.org/crap/). Peptide and protein identifications were validated using Scaffold (v5.3.3, www.proteomesoftware.com), which was also used to calculate normalized spectral abundance factors (NSAFs). Proteins were required to have a minimum of two peptides and meet false discovery rate (FDR) thresholds of 0.1% at the peptide level and 1% at the protein level. To ensure comparability across samples, the two lowest-yielding heavyIE and lightThy samples (based on total spectral counts) were excluded from downstream analysis. The final dataset consisted of 19 samples: four inner envelope (IE), five heavyIE, four thylakoid (Thy), and six light thylakoid (lightThy) replicates. Data were processed in Perseus v2.1.3.0 (Rudolph and Cox, 2019). NSAF values were log2- transformed, and known contaminants were removed. Protein identifications were filtered to retain only those detected in at least three biological replicates within a single sample type and at least four replicates total. Missing values were imputed from a normal distribution. Statistical comparisons were performed using a two-sided t-test modified for FDR control by permutation (Tyanova et al., 2016). Proteins were considered significantly differentially abundant if they met a log2 fold change threshold of ±0.5 and an adjusted p-value cutoff of –log10(p) 2: 1.15 (FDR :5 5%). Subcellular localization of identified proteins was inferred from their Arabidopsis homologs using manually integrated data from SUBA (Hooper et al., 2014) and PPDB (Sun et al., 2009). Principal Component Analysis was performed in R using prcomp (Team, 2023)(R Core Team, 2023), and volcano plots and KEGG pathway enrichment were visualized with ggplot2 (Wickham, 2009). To assess similarity to known membrane contact site (MCS) proteins, sequences of Arabidopsis homolog proteins were queried against the MCSdatabase v 1.7 (Pan et al., 2024). Proteins with bit scores over 50 were considered to share homology with MCS components.

### Transmission Electron Microscopy and Analysis

Seedlings (27 h post-germination) were fixed, embedded, sectioned, post-stained, and imaged as described (Keynia et al., 2023), except fix occurred under 1.5 mTorr vacuum for 1 h, and thin sections were 70 nm. TIES were defined as thylakoid–inner envelope contacts :5 50 nm apart. Length and intermembrane distances were measured using FIJI (Schindelin et al., 2012) with the InteredgeDistance macro (version 1.3, Santosh Patnaik). Chloroplast area was measured with FIJI. Statistical analysis was performed in GraphPad Prism using one-way ANOVA followed by Dunnett’s corrected post hoc tests.

### Structural Predictions

Predicted structures of TVPFP and PMFP were generated using AlphaFold (Jumper et al., 2021; Varadi et al., 2022) after which membrane binding regions were predicted using PPM 3.0 (Lomize et al., 2022).

## Acknowledgements

The authors gratefully acknowledge Samantha Link and Kandy Hanthorn for excellent plant management, Dr. You Zhou for management of the Microscopy Center (RRID:SCR 017798) and Dr. Sophie Alvarez for management of the Proteomics and Metabolomics Facility (RRID:SCR_021314). This project was supported by the Department of Energy (DE-SC0021101) and partially supported by the Nebraska Agricultural Experiment Station with funding from the Hatch Multistate capacity funding program (Accession Number 7003614) from the USDA National Institute of Food and Agriculture.

## Author Contributions

EWL drafted the original manuscript. EWL, CNS, and RLR conceptualized the project. EWL, CNS, AT-G, JI, BA, MJN, and RLR contributed to investigation. Data curation, validation, and visualization were performed by EWL, CNS, AT-G, JI, F-H, and RLR. Formal analysis was conducted by EWL, CNS, AT-G, JI, F-H, AT, LL, BM, and RLR. BA and RLR supervised the project. RLR provided funding acquisition, project administration, and resources. All authors reviewed and edited the final manuscript.

## Supplementary Figures

**Figure S1.**
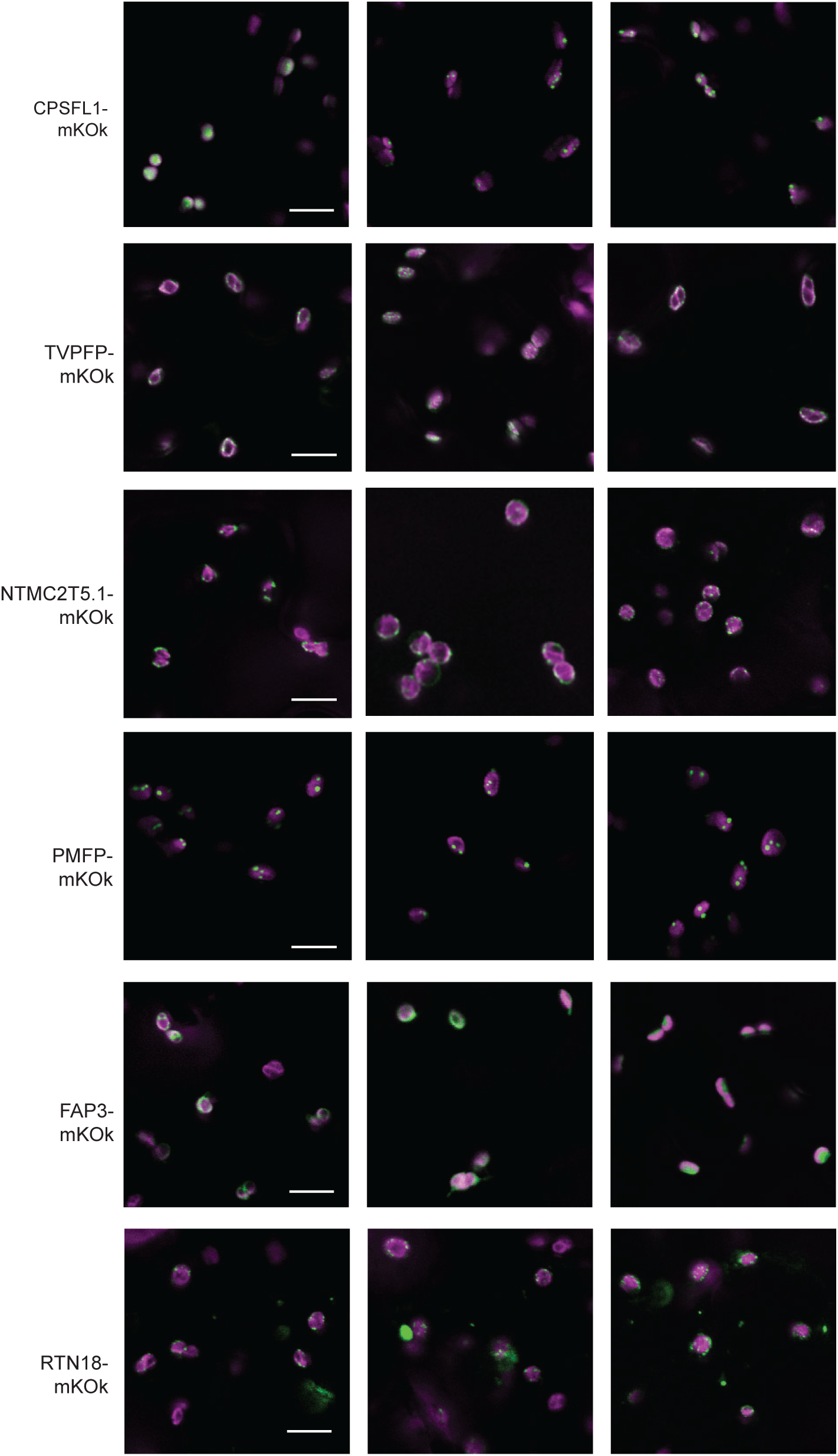
Additional Confocal Images of Chloroplast Localized Candidate Proteins. Representative confocal images of mKOK-tagged candidates expressed in *N. benthamiana*. TVPFP, NTMC2T5.1, PMFP, and RTN18 displayed punctate or reticulate fluorescence patterns suggestive of functionally specialized chloroplast regions. FAP3 exhibited a diffuse chloroplast signal. CPSFL1 showed both diffuse and punctate fluorescence, consistently overlapping with chlorophyll. Scale bar = 10 μm. Abbreviations: mKOK, monomeric Kusabira Orange kappa; CPSFL1, Chloroplast Sec14-like 1; TVPFP, Tvp38 Family Protein; NTMC2T5.1, N-Terminal Transmembrane C2 Domain Type 5.1; PMFP, Plasma Membrane Fusion Protein; FAP3, Fatty Acid Binding Protein 3; RTN18, Reticulon 18.

**Figure S2:**
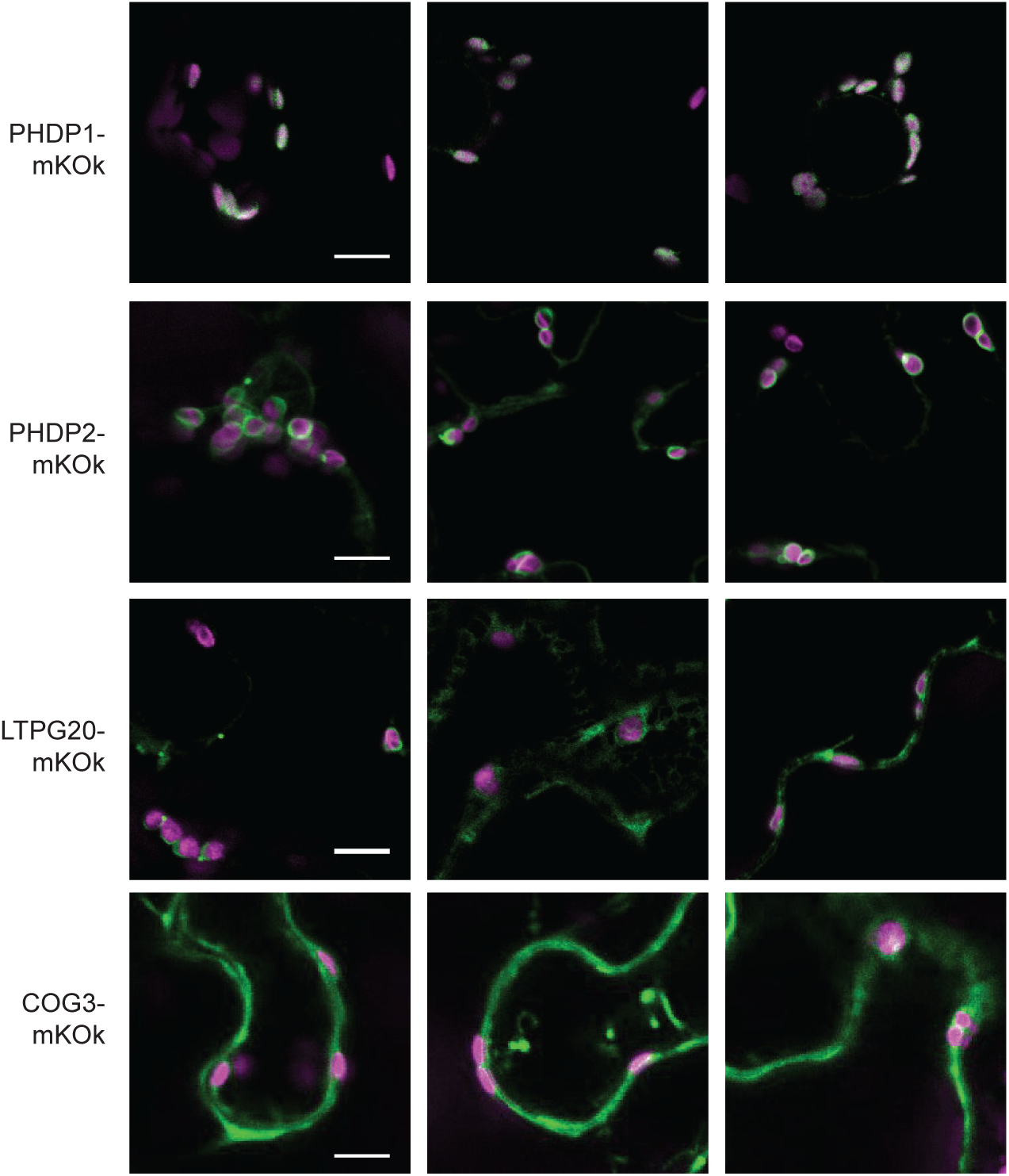
Additional Confocal Images of Candidates Showing Mixed Localization. Representative confocal images of mKOK-tagged proteins PHDP1, PHDP2, LTPG20, and COG3. PHDP1/2 showed strong but non-exclusive chloroplast association. LTPG20 and COG3 exhibited overlapping but also non-chloroplast localization. Scale bar = 10 μm. Abbreviations: mKOK, monomeric Kusabira Orange kappa; PHDP1/2, Pleckstrin Homology Domain Protein; LTPG20, GLYCOSYLPHOSPHATIDYLINOSITOL- ANCHORED LIPID PROTEIN TRANSER 20; COG3, Conserved Oligomeric Golgi complex protein 3.

**Figure S3:**
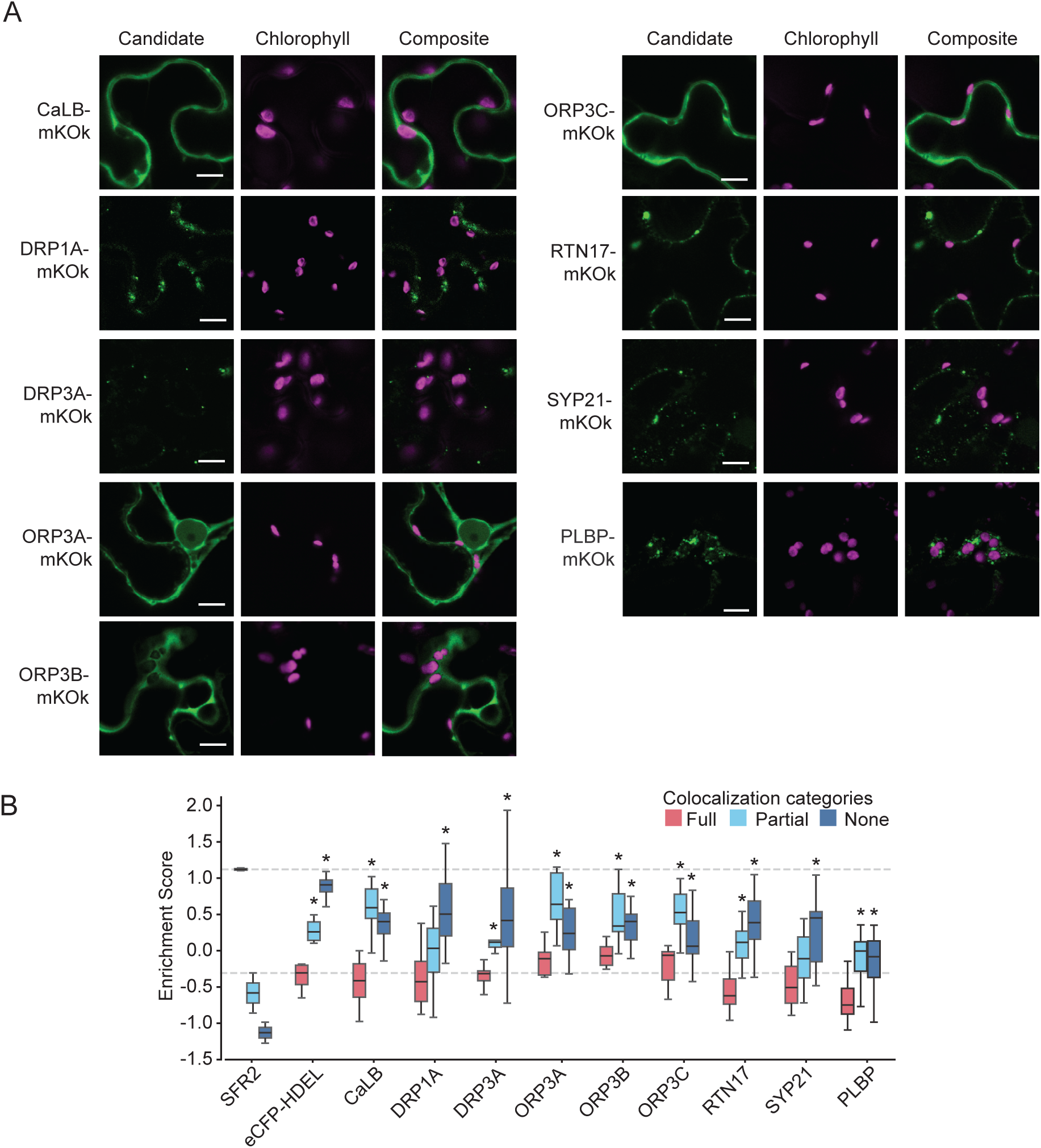
Candidate proteins without detectable chloroplast colocalization. (A) Representative confocal images of mKOK tagged candidate proteins showing little to no overlap with chlorophyll autofluorescence. Insets = 4 × 4 μm. Scale bar = 10 μm. (B) CellProfiler-based quantification of colocalization between candidate signals and chlorophyll autofluorescence. Colocalization categories are represented as boxplots (n 2: 3 trials, >100 chloroplasts total). Dashed lines indicate the median values for full colocalization of the positive (SFR2-eGFP) and negative (eCFP-HDEL) controls. Asterisks denote statistical significance (*p* < 0.05) by Welch’s ANOVA followed by Dunnett’s T3 multiple comparisons test between the full colocalization group and other categories for each protein (see Table S2 for full statistics). Abbreviations: mKOK, monomeric Kusabira Orange kappa; CaLB, Calcium-dependent Lipid Binding protein; DRP1A, Dynamin-Related Protein 1A; DRP3A, Dynamin-Related Protein 3A; ORP3A/B/C, Oxysterol binding Protein 3A/B/C; RTN17, Reticulon 17; SYP21, Syntaxin in Plants 21; PLBP, Putative Lipid Binding Protein

**Figure S4.**
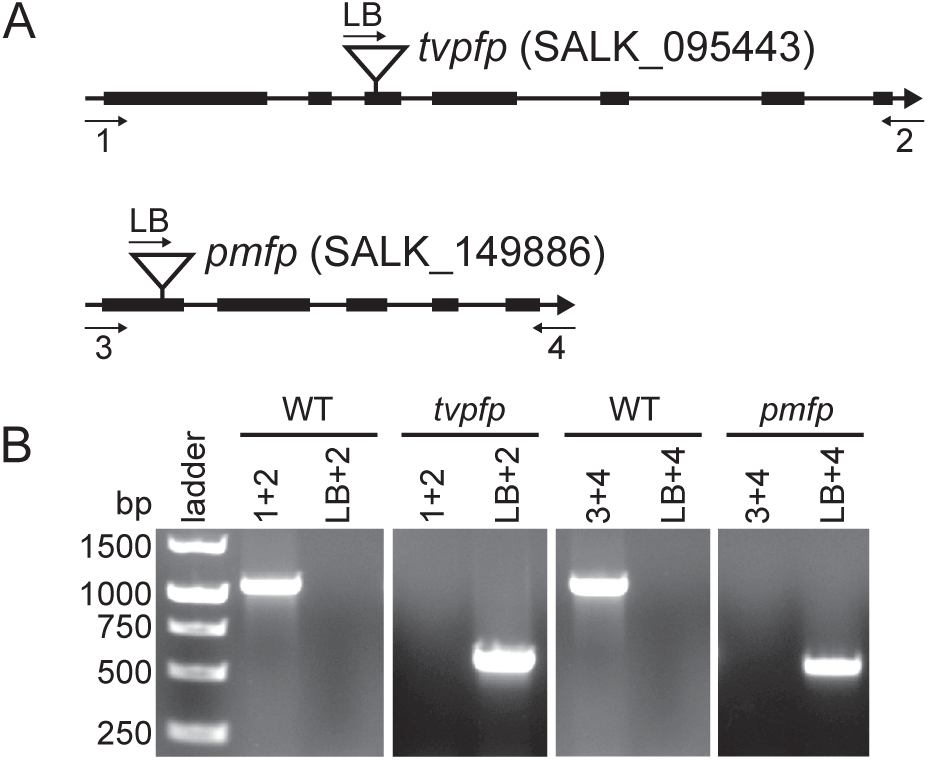
PCR Verification of *tvpfp* and *pmfp* T-DNA Insertional Mutants. (A) Gene models showing exon/intron structure and T-DNA insertion sites for *TVPFP* and *PMFP*. Primer binding sites are indicated. (B) PCR results for mutant genotyping. Homozygous T-DNA insertions produced expected band sizes and lacked WT-specific bands for both *tvpfp* (SALK_095443) and *pmfp* (SALK_149886). Abbreviations: WT, wildtype; TVPFP, TVP38 FAMILY PROTEIN; PMFP, PLASMA MEMBRANE FUSION PROTEIN.

**Figure S5.**
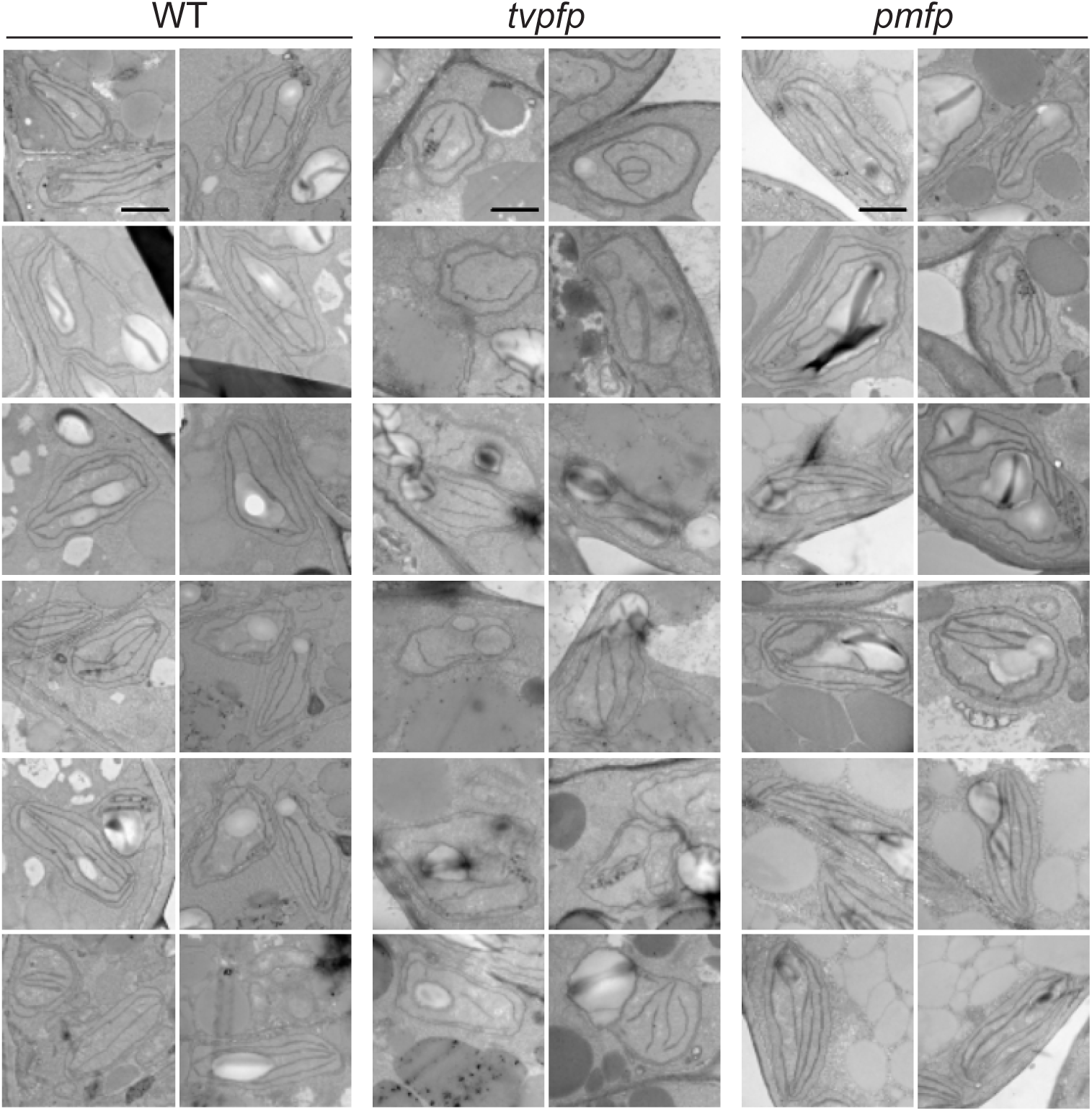
Additional Transmission Electron Micrographs of *tvpfp* and *pmfp* Chloroplasts. Transmission electron micrographs of chloroplasts from WT, *tvpfp*, and *pmfp* seedlings at 27 hours after germination. WT and *pmfp* plastids display typical lenticular shape and polar thylakoid-envelope proximity. *tvpfp* mutants show irregular shapes and disrupted thylakoid organization. Scale bar = 1 μm.

## Supplementary Tables

Table S1. **Candidate Proteins with Potential Roles in Chloroplast Lipid Transport or Membrane Contact Sites.** Candidate proteins investigated for chloroplast localization and possible involvement in lipid transfer or membrane contact sites.

Table S2. **Statistical summary of CellProfiler fluorescence colocalization analysis.** Welch’s ANOVA with Dunnett’s T3 multiple comparisons test results for fluorescent protein signal colocalization across candidate genes. Comparisons include full, partial, and no overlap with chlorophyll autofluorescence.

Table S3. **Proteins identified by LC-MS/MS across chloroplast membrane fractions.**

List of all proteins detected in the proteomic analysis with corresponding Arabidopsis gene identifiers, peptide counts, sequence coverage, and confidence scores.

Table S4. **Detected protein normalized abundance values, fraction enrichment, and subcellular localization annotation.**

LFQ intensity values for each protein across biological replicates of inner envelope (IE), heavy inner envelope (heavyIE), thylakoid (Thy), and light thylakoid (lightThy) membrane fractions. Proteins showing significant enrichment in pairwise comparisons between membrane fractions are indicated. Includes fold changes, p-values, and enrichment direction. Subcellular and suborganellar localization of all identified proteins based on curated annotations from SUBA, PPDB, and literature sources.

Table S5. **Proteins with homology to membrane contact site (MCS) components.** Proteins enriched in each fraction with sequence similarity to known MCS-associated proteins.

Table S6. **Pathway enrichment results.**

Functional enrichment output from KEGG and GO analysis of proteins enriched in the all fractions, including pathway name, size, FDR, and protein representation.

Table S7. **Perturbation of Chloroplast Ultrastructure by Loss of TVPFP or PMFP.** Ultrastructural measurements from transmission electron micrographs of palisade mesophyll chloroplasts in wild-type (WT), *tvpfp*, and *pmfp* seedlings.

## REFERENCES

Alonso, J.M., Stepanova, A.N., Leisse, T.J., Kim, C.J., Chen, H., Shinn, P., Stevenson, D.K., Zimmerman, J., Barajas, P., Cheuk, R., et al. (2003). Genome-wide insertional mutagenesis of Arabidopsis thaliana. Science 301, 653–657.

Bouchnak, I., Brugière, S., Moyet, L., Le Gall, S., Salvi, D., Kuntz, M., Tardif, M., and Rolland, N. (2019). Unravelling hidden components of the chloroplast envelope proteome: opportunities and limits of better MS sensitivity. Molecular &amp; Cellular Proteomics, mcp.RA118.000988.

Boudière, L., Michaud, M., Petroutsos, D., Rébeillé, F., Falconet, D., Bastien, O., Roy, S., Finazzi, G., Rolland, N., Jouhet, J., et al. (2014). Glycerolipids in photosynthesis: Composition, synthesis and trafficking. Biochimica et Biophysica Acta (BBA) - Bioenergetics 1837, 470–480.

Boughner, L.A., and Doerrler, W.T. (2012). Multiple deletions reveal the essentiality of the DedA membrane protein family in Escherichia coli. Microbiology 158, 1162–1171.

Bruce, B.D., Perry, S., Froehlich, J., and Keegstra, K. (1994). In vitro import of proteins into chloroplasts. In Plant Molecular Biology Manual, S.B. Gelvin, and R.A. Schilperoort, eds. (Dordrecht: Springer Netherlands), pp. 497–511.

Cackett, L., Luginbuehl, L.H., Schreier, T.B., Lopez-Juez, E., and Hibberd, J.M. (2022). Chloroplast development in green plant tissues: The interplay between light, hormone, and transcriptional regulation. New Phytol 233, 2000–2016.

Cai, Y., Goodman, J.M., Pyc, M., Mullen, R.T., Dyer, J.M., and Chapman, K.D. (2015). Arabidopsis SEIPIN Proteins Modulate Triacylglycerol Accumulation and Influence Lipid Droplet Proliferation. The Plant Cell 27, 2616–2636.

Charuvi, D., Kiss, V., Nevo, R., Shimoni, E., Adam, Z., and Reich, Z. (2012). Gain and Loss of Photosynthetic Membranes during Plastid Differentiation in the Shoot Apex of Arabidopsis. Plant Cell 24, 1143–1157.

Cook, R., Lupette, J., and Benning, C. (2021). The Role of Chloroplast Membrane Lipid Metabolism in Plant Environmental Responses. Cells 10.

Dorne, A.J., Joyard, J., and Douce, R. (1990). Do thylakoids really contain phosphatidylcholine? ProcNatlAcadSciUSA 87, 71–74.

Douce, R. (1974). Site of biosynthesis of galactolipids in spinach chloroplasts. Science 183, 852–853.

Douce, R., and Joyard, J. (1996). Biosynthesis of thylakoid membrane lipids. In Oxygenic Photosynthesis: The Light Reactions, D.R. Ort, and C.F. Yocum, eds. (Dordrecht: Kluwer Academic Publishers), pp. 69-101.

Fan, J., Zhai, Z., Yan, C., and Xu, C. (2015). Arabidopsis TRIGALACTOSYLDIACYLGLYCEROL5 Interacts with TGD1, TGD2, and TGD4 to Facilitate Lipid Transfer from the Endoplasmic Reticulum to Plastids. Plant Cell 27, 2941–2955.

Ferro, M., Brugiere, S., Salvi, D., Seigneurin-Berny, D., Court, M., Moyet, L., Ramus, C., Miras, S., Mellal, M., Le Gall, S., et al. (2010). AT_CHLORO, a Comprehensive Chloroplast Proteome Database with Subplastidial Localization and Curated Information on Envelope Proteins. Mol Cell Proteomics 9, 1063–1084.

Ferro, M., Salvi, D., Brugière, S., Miras, S., Kowalski, S., Louwagie, M., Garin, J., Joyard, J., and Rolland, N. (2003). Proteomics of the chloroplast envelope membranes from Arabidopsis thaliana. Molecular & cellular proteomics: MCP 2, 325–345.

Fujii, S., Nagata, N., Masuda, T., Wada, H., and Kobayashi, K. (2019). Galactolipids Are Essential for Internal Membrane Transformation during Etioplast-to-Chloroplast Differentiation. Plant Cell Physiol 60, 1224–1238.

García-Cerdán, J.G., Schmid, E.M., Takeuchi, T., McRae, I., McDonald, K.L., Yordduangjun, N., Hassan, A.M., Grob, P., Xu, C.S., Hess, H.F., et al. (2020). Chloroplast Sec14-like 1 (CPSFL1) is essential for normal chloroplast development and affects carotenoid accumulation in Chlamydomonas. Proceedings of the National Academy of Sciences 117, 12452–12463.

Ghanbarpour, A., Valverde, D.P., Melia, T.J., and Reinisch, K.M. (2021). A model for a partnership of lipid transfer proteins and scramblases in membrane expansion and organelle biogenesis. P Natl Acad Sci USA 118, 2101562118.

Gibbs, S.P. (1962). The ultrastructure of the chloroplasts of algae. Journal of Ultrastructure Research 7, 418–435.

Gounaris, K., Mannock, D.D., Sen, A., Brain, A.P.R., Wiliams, W.P., and Quinn, P.J. (1983). Polyunsaturated fatty acyl residues of galactolipids are involved in the control of bilayer/nonbilayer lipid transitions in higher plant chloroplasts. BiochimBiophysActa 732, 229–242.

Gupta, T.K., Klumpe, S., Gries, K., Heinz, S., Wietrzynski, W., Ohnishi, N., Niemeyer, J., Spaniol, B., Schaffer, M., Rast, A., et al. (2021). Structural basis for VIPP1 oligomerization and maintenance of thylakoid membrane integrity. Cell 184, 3643–3659 3623.

Hermann, G.J., Thatcher, J.W., Mills, J.P., Hales, K.G., Fuller, M.T., Nunnari, J., and Shaw, J.M. (1998). Mitochondrial Fusion in Yeast Requires the Transmembrane GTPase Fzo1p. J Cell Biol 143, 359–373.

Hertle, A.P., García-Cerdán, J.G., Armbruster, U., Shih, R., Lee, J.J., Wong, W., and Niyogi, K.K. (2020). A Sec14 domain protein is required for photoautotrophic growth and chloroplast vesicle formation in Arabidopsis thaliana. Proceedings of the National Academy of Sciences 117, 9101–9111.

Hölzl, G., and Dörmann, P. (2019). Chloroplast lipids and their biosynthesis. Annu Rev Plant Biol 70, 51–81.

Hooper, C., Millar, H., Black, K., Castleden, I., Aryamanesh, N., and Grasso, S. (2022). Subcellular Localisation database for Arabidopsis proteins version 5. The UWA Profiles and Research Repository.

Hooper, C.M., Tanz, S.K., Castleden, I.R., Vacher, M.A., Small, I.D., and Millar, A.H. (2014). SUBAcon: a consensus algorithm for unifying the subcellular localization data of the Arabidopsis proteome. Bioinformatics 30, 3356–3364.

Huang, D., Xu, B., Liu, L., Wu, L., Zhu, Y., Ghanbarpour, A., Wang, Y., Chen, F.-J., Lyu, J., Hu, Y., et al. (2021). TMEM41B acts as an ER scramblase required for lipoprotein biogenesis and lipid homeostasis. Cell Metabolism 33, 1655–1670 1658.

Huercano, C., Moya-Barrientos, M., Cuevas, O., Sanchez-Vera, V., and Ruiz-Lopez, N. (2025). ER–plastid contact sites as molecular crossroads for plastid lipid biosynthesis. BMC Biol 23, 139.

Hunter, J.D. (2007). Matplotlib: A 2D graphics environment. Computing in science & engineering 9, 90–95.

Ishidate, K., Creeger, E.S., Zrike, J., Deb, S., Glauner, B., MacAlister, T.J., and Rothfield, L.I. (1986). Isolation of differentiated membrane domains from Escherichia coli and Salmonella typhimurium, including a fraction containing attachment sites between the inner and outer membranes and the murein skeleton of the cell envelope. J Biol Chem 261, 428–443.

Jarvis, P., and López-Juez, E. (2013). Biogenesis and homeostasis of chloroplasts and other plastids. Nature Reviews Molecular Cell Biology 14, 787–802.

Jouhet, J., Gros, V., and Michaud, M. (2024). Measurement of Lipid Transport in Mitochondria by the MTL Complex. In Intracellular Lipid Transport: Methods and Protocols (Springer), pp. 167–191.

Jumper, J., Evans, R., Pritzel, A., Green, T., Figurnov, M., Ronneberger, O., Tunyasuvunakool, K., Bates, R., Zidek, A., Potapenko, A., et al. (2021). Highly accurate protein structure prediction with AlphaFold. Nature 596, 583–589.

Junglas, B., Kartte, D., Kutzner, M., Hellmann, N., Ritter, I., Schneider, D., and Sachse, C. (2025). Structural basis for Vipp1 membrane binding: From loose coats and carpets to ring and rod assemblies. Nat Struct Mol Biol 32, 555–570.

Keegstra, K., and Yousif, A.E. (1986). Isolation and characterization of chloroplast envelope membranes. In Methods Enzymol (Academic Press), pp. 316–325.

Keynia, S., Jaafar, L., Zhou, Y., Anderson, C.T., and Turner, J.A. (2023). Stomatal opening efficiency is controlled by cell wall organization in Arabidopsis thaliana. PNAS Nexus 2.

Kim, E., Poudyal, R.S., Lee, K., Yu, H., Gi, E., and Kim, H.U. (2022). Chloroplast- localized PITP7 is essential for plant growth and photosynthetic function in Arabidopsis. Physiol Plant 174, 13760.

Krol, M., Huner, N.P.A., and McIntosh, A. (1987). Chloroplast biogenesis at cold-hardening temperatures. Development of photosystem I and photosystem II activities in relation to pigment accumulation. Photosynthesis Res 14, 97–112.

Kroll, D., Meierhoff, K., Bechtold, N., Kinoshita, M., Westphal, S., Vothknecht, U.C., Soll, J., and Westhoff, P. (2001). VIPP1, a nuclear gene of Arabidopsis thaliana essential for thylakoid membrane formation. P Natl Acad Sci USA 98, 4238–4242.

LaBrant, E., Barnes, A.C., and Roston, R.L. (2018). Lipid transport required to make lipids of photosynthetic membranes. Photosynth Res 138, 345–360.

Lahiri, S., Toulmay, A., and Prinz, W.A. (2015). Membrane contact sites, gateways for lipid homeostasis. Curr Opin Cell Biol 33, 82–87.

Li-Beisson, Y., Shorrosh, B., Beisson, F., Andersson, M.X., Arondel, V., Bates, P.D., Baud, S., Bird, D., Debono, A., Durrett, T.P., et al. (2013). Acyl-lipid metabolism. Arabidopsis Book 11, e0161.

Li, Y.E., Wang, Y., Du, X., Zhang, T., Mak, H.Y., Hancock, S.E., McEwen, H., Pandzic, E., Whan, R.M., Aw, Y.C., et al. (2021). TMEM41B and VMP1 are scramblases and regulate the distribution of cholesterol and phosphatidylserine. The Journal of Cell Biology 220, 202103105.

Liang, Z., Zhu, N., Mai, K.K., Liu, Z., Tzeng, D., Osteryoung, K.W., Zhong, S., Staehelin, L.A., and Kang, B.-H. (2018). Thylakoid-Bound Polysomes and a Dynamin-Related Protein, FZL, Mediate Critical Stages of the Linear Chloroplast Biogenesis Program in Greening Arabidopsis Cotyledons. The Plant Cell 30, 1476–1495.

Lin, J.J. (1995). Electrotransformation of Agrobacterium. Methods in Molecular Biology.

Lindquist, E., Solymosi, K., and Aronsson, H. (2016). Vesicles Are Persistent Features of Different Plastids. Traffic 17, 1125–1138.

Lomize, A.L., Todd, S.C., and Pogozheva, I.D. (2022). Spatial arrangement of proteins in planar and curved membranes by PPM 3.0. Protein Sci 31, 209–220.

Loudya, N., Barkan, A., and López-Juez, E. (2024). Plastid retrograde signaling: A developmental perspective. The Plant Cell 36, 3903–3913.

Lu, B., Xu, C., Awai, K., Jones, A.D., and Benning, C. (2007). A small ATPase protein of Arabidopsis, TGD3, involved in chloroplast lipid import. J Biol Chem 282, 35945–35953.

Mai, K.K.K., Yeung, W.-T., Han, S.-Y., Cai, X., Hwang, I., and Kang, B.-H. (2019). Electron Tomography Analysis of Thylakoid Assembly and Fission in Chloroplasts of a Single-Cell C4 plant, Bienertia sinuspersici. Sci Rep-Uk 9, 19640.

Mares, J., Strunecky, 0., Bucinska, L., and Wiedermannova, J. (2019). Evolutionary Patterns of Thylakoid Architecture in Cyanobacteria. Front Microbiol 10.

Mazur, R., Gieczewska, K., Kowalewska, t., Kuta, A., Proboszcz, M., Gruszecki, W.I., Mostowska, A., and Garstka, M. (2020). Specific Composition of Lipid Phases Allows Retaining an Optimal Thylakoid Membrane Fluidity in Plant Response to Low-Temperature Treatment. Frontiers in Plant Science 11.

McKinney, W. (2011). pandas: a foundational Python library for data analysis and statistics. Python for high performance and scientific computing 14, 1–9.

Mi, H., Ebert, D., Muruganujan, A., Mills, C., Albou, L.-P., Mushayamaha, T., and Thomas, P.D. (2021). PANTHER version 16: A revised family classification, tree-based classification tool, enhancer regions and extensive API. Nucleic Acids Res 49, 394– 403.

Morré, D.J., Sellden, G., Sundqvist, C., and Sandelius, A.S. (1991). Stromal Low Temperature Compartment Derived from the Inner Membrane of the Chloroplast Envelope. Plant Physiol 97, 1558–1564.

Nakamura, S., Matsui, A., Akabane, S., Tamura, Y., Hatano, A., Miyano, Y., Omote, H., Kajikawa, M., Maenaka, K., Moriyama, Y., et al. (2020). The mitochondrial inner membrane protein LETM1 modulates cristae organization through its LETM domain. Communications Biology 3, 1–11.

Norkunas, K., Harding, R., Dale, J., and Dugdale, B. (2018). Improving agroinfiltration-based transient gene expression in Nicotiana benthamiana. Plant Methods 14, 71.

Ostermeier, M., Garibay-Hernández, A., Holzer, V.J., Schroda, M., and Nickelsen, J. (2024). Structure, biogenesis, and evolution of thylakoid membranes. The Plant Cell 36, 4014–4035.

Páli, T., Garab, G., Horváth, L.I., and Kóta, Z. (2003). Functional significance of the lipid-protein interface in photosynthetic membranes. Cellular and Molecular Life Sciences: CMLS 60, 1591–1606.

Pan, X., Ren, L., Yang, Y., Xu, Y., Ning, L., Zhang, Y., Luo, H., Zou, Q., and Zhang, Y. (2024). MCSdb, a database of proteins residing in membrane contact sites. Scientific Data 11, 281.

Peltier, J.B., Ytterberg, A.J., Sun, Q., and van Wijk, K.J. (2004). New functions of the thylakoid membrane proteome of Arabidopsis thaliana revealed by a simple, fast, and versatile fractionation strategy. J Biol Chem 279, 49367–49383.

Petroutsos, D., Amiar, S., Abida, H., Dolch, L.-J., Bastien, O., Rébeillé, F., Jouhet, J., Falconet, D., Block, M.A., McFadden, G.I., et al. (2014). Evolution of galactoglycerolipid biosynthetic pathways – From cyanobacteria to primary plastids and from primary to secondary plastids. Prog Lipid Res 54, 68–85.

Pineau, B., Gerardhirne, C., Douce, R., and Joyard, J. (1993). Identification of the Main Species of Tetrapyrrolic Pigments in Envelope Membranes from Spinach-Chloroplasts. Plant Physiol 102, 821–828.

Pon, L., Moll, T., Vestweber, D., Marshallsay, B., and Schatz, G. (1989). Protein import into mitochondria: ATP-dependent protein translocation activity in a submitochondrial fraction enriched in membrane contact sites and specific proteins. The Journal of Cell Biology 109, 2603–2616.

Quon, E., and Beh, C.T. (2015). Membrane Contact Sites: Complex Zones for Membrane Association and Lipid Exchange. Lipid Insights 8, 55–63.

Rast, A., Schaffer, M., Albert, S., Wan, W., Pfeffer, S., Beck, F., Plitzko, J.M., Nickelsen, J., and Engel, B.D. (2019). Biogenic regions of cyanobacterial thylakoids form contact sites with the plasma membrane. Nature Plants 5, 436–446.

Rawyler, A., Meylan, M., and Siegenthaler, P.A. (1992). Galactolipid export from envelope to thylakoid membranes in intact chloroplasts. I. Characterization and involvement in thylakoid lipid asymmetry. Biochimica Biophysica Acta 1104, 331–341.

Roston, R.L., Gao, J.P., Murcha, M.W., Whelan, J., and Benning, C. (2012). TGD1,-2, and-3 proteins involved in lipid trafficking form ATP-binding cassette (ABC) transporter with multiple substrate-binding proteins. J Biol Chem 287, 21406–21415.

Roston, R.L., Wang, K., Kuhn, L.A., and Benning, C. (2014). Structural determinants allowing transferase activity in SENSITIVE TO FREEZING 2, classified as a family I glycosyl hydrolase. J Biol Chem 289, 26089–26106.

Rudolph, J.D., and Cox, J. (2019). A Network Module for the Perseus Software for Computational Proteomics Facilitates Proteome Interaction Graph Analysis. J Proteome Res 18, 2052–2064.

Rusiñol, A.E., Cui, Z., Chen, M.H., and Vance, J.E. (1994). A unique mitochondria-associated membrane fraction from rat liver has a high capacity for lipid synthesis and contains pre-Golgi secretory proteins including nascent lipoproteins. J Biol Chem 269, 27494–27502.

Sato, N., and Awai, K. (2016). Diversity in Biosynthetic Pathways of Galactolipids in the Light of Endosymbiotic Origin of Chloroplasts. Frontiers in Plant Science 7.

Schindelin, J., Arganda-Carreras, I., Frise, E., Kaynig, V., Longair, M., Pietzsch, T., Preibisch, S., Rueden, C., Saalfeld, S., Schmid, B., et al. (2012). Fiji: An open-source platform for biological-image analysis. Nat Methods 9, 676–682.

Schneider, C.A., Rasband, W.S., and Eliceiri, K.W. (2012). NIH Image to ImageJ: 25 years of image analysis. Nat Methods 9, 671–675.

Seifert, U., and Heinz, E. (1992). Enzymatic characteristics of UDP-sulfoquinovose:diacylglycerol sulfoquinovosyltranferase from chloroplast envelopes. BotActa 105, 197–205.

Shimoni, E., Rav-Hon, O., Ohad, I., Brumfeld, V., and Reich, Z. (2005). Three-Dimensional Organization of Higher-Plant Chloroplast Thylakoid Membranes Revealed by Electron Tomography. THE PLANT CELL 17, 2580–2586.

Shivaiah, K.K., Susanto, F.A., Devadasu, E., and Lundquist, P.K. (2022). Plastoglobule Lipid Droplet Isolation from Plant Leaf Tissue and Cyanobacteria. J Vis Exp.

Shomo, Z.D., Mahboub, S., Vanviratikul, H., McCormick, M., Tulyananda, T., Roston, R.L., and Warakanont, J. (2024). All members of the Arabidopsis DGAT and PDAT acyltransferase families operate during high and low temperatures. Plant Physiol 195, 685–697.

Soll, J., Schultz, G., Rüdiger, W., and Benz, J. (1983). Hydrogenation of Geranylgeraniol 1: Two Pathways Exist in Spinach Chloroplasts. Plant Physiol 71, 849–854.

Staehelin, L.A. (1986). Chloroplast Structure and Supramolecular Organization of Photosynthetic Membranes. In Photosynthesis III: Photosynthetic Membranes and Light Harvesting Systems, L.A. Staehelin, and C.J. Arntzen, eds. (Springer), pp. 1–84.

Stirling, D.R., Carpenter, A.E., and Cimini, B.A. (2021a). CellProfiler Analyst 3.0: Accessible data exploration and machine learning for image analysis. Bioinformatics 37, 3992–3994.

Stirling, D.R., Swain-Bowden, M.J., Lucas, A.M., Carpenter, A.E., Cimini, B.A., and Goodman, A. (2021b). CellProfiler 4: Improvements in speed, utility and usability. BMC Bioinformatics 22, 433.

Stone, A.B. (1974). A simplified method for preparing sucrose gradients. Biochem J 137, 117–118.

Sun, Q., Zybailov, B., Majeran, W., Friso, G., Olinares, P.D.B., and van Wijk, K.J. (2009). PPDB, the Plant Proteomics Database at Cornell. Nucleic Acids Res 37, D969–D974.

Sun, Y., Bakhtiari, S., Valente-Paterno, M., Jiang, H., and Zerges, W. (2025). Membranous translation platforms in the chloroplast of Chlamydomonas reinhardtii. Plant Physiol 197.

Tábara, L.-C., and Escalante, R. (2016). VMP1 Establishes ER-Microdomains that Regulate Membrane Contact Sites and Autophagy. PLoS One 11, 0166499.

Team, R.C. (2023). R: A Language and Environment for Statistical Computing. R Foundation for Statistical Computing, Vienna.

Tefsen, B., Geurtsen, J., Beckers, F., Tommassen, J., and de Cock, H. (2005). Lipopolysaccharide transport to the bacterial outer membrane in spheroplasts. JBiolChem 280, 4504–4509.

Tomizioli, M., Lazar, C., Brugière, S., Burger, T., Salvi, D., Gatto, L., Moyet, L., Breckels, L.M., Hesse, A.-M., Lilley, K.S., et al. (2014). Deciphering Thylakoid Sub-compartments using a Mass Spectrometry-based Approach *. Mol Cell Proteomics 13, 2147–2167.

Trotta, A., Bajwa, A.A., Mancini, I., Paakkarinen, V., Pribil, M., and Aro, E.-M. (2019). The Role of Phosphorylation Dynamics of CURVATURE THYLAKOID 1B in Plant Thylakoid Membranes. Plant Physiol 181, 1615–1631.

Tsutsui, H., Karasawa, S., Okamura, Y., and Miyawaki, A. (2008). Improving membrane voltage measurements using FRET with new fluorescent proteins. Nat Methods 5, 683–685.

Tyanova, S., Temu, T., Sinitcyn, P., Carlson, A., Hein, M.Y., Geiger, T., Mann, M., and Cox, J. (2016). The Perseus computational platform for comprehensive analysis of (prote)omics data. Nat Methods 13, 731–740.

Vance, J.E. (2014). MAM (mitochondria-associated membranes) in mammalian cells: Lipids and beyond. Biochimica et Biophysica Acta (BBA) - Molecular and Cell Biology of Lipids 1841, 595–609.

Varadi, M., Anyango, S., Deshpande, M., Nair, S., Natassia, C., Yordanova, G., Yuan, D., Stroe, O., Wood, G., Laydon, A., et al. (2022). AlphaFold Protein Structure Database: massively expanding the structural coverage of protein-sequence space with high-accuracy models. Nucleic Acids Res 50, D439–D444.

Wang, L., Patena, W., Van Baalen, K.A., Xie, Y., Singer, E.R., Gavrilenko, S., Warren-Williams, M., Han, L., Harrigan, H.R., Hartz, L.D., et al. (2023). A chloroplast protein atlas reveals punctate structures and spatial organization of biosynthetic pathways. Cell 186, 3499–3518.e3414.

Waskom, M.L. (2021). Seaborn: statistical data visualization. Journal of Open Source Software 6, 3021.

Weiss, G.L., Eisenstein, F., Kieninger, A.-K., Xu, J., Minas, H.A., Gerber, M., Feldmüller, M., Maldener, I., Forchhammer, K., and Pilhofer, M. (2022). Structure of a thylakoid-anchored contractile injection system in multicellular cyanobacteria. Nature Microbiology 7, 386–396.

Wickham, H. (2009). ggplot2: Elegant graphics for data analysis. Springer-Verlag, NY.

Wietrzynski, W., Schaffer, M., Tegunov, D., Albert, S., Kanazawa, A., Plitzko, J.M., Baumeister, W., and Engel, B.D. (2020). Charting the native architecture of Chlamydomonas thylakoid membranes with single-molecule precision. eLife 9, 53740.

Yoshihara, A., Nagata, N., Wada, H., and Kobayashi, K. (2021). Plastid Anionic Lipids Are Essential for the Development of Both Photosynthetic and Non-Photosynthetic Organs in Arabidopsis thaliana. Int J Mol Sci 22.

Zhang, L., Kato, Y., Otters, S., Vothknecht, U.C., and Sakamoto, W. (2012). Essential Role of VIPP1 in Chloroplast Envelope Maintenance in Arabidopsis. The Plant Cell 24, 3695–3707.

Zhao, Y.G., Chen, Y., Miao, G., Zhao, H., Qu, W., Li, D., Wang, Z., Liu, N., Li, L., Chen, S., et al. (2017). The ER-Localized Transmembrane Protein EPG-3/VMP1 Regulates SERCA Activity to Control ER-Isolation Membrane Contacts for Autophagosome Formation. Mol Cell 67, 974–989 976.

Zhu, D., Xiong, H., Wu, J., Zheng, C., Lu, D., Zhang, L., and Xu, X. (2022). Protein Targeting Into the Thylakoid Membrane Through Different Pathways. Front Physiol 12, 802057.

